# Analysis of multi-condition single-cell data with latent embedding multivariate regression

**DOI:** 10.1101/2023.03.06.531268

**Authors:** Constantin Ahlmann-Eltze, Wolfgang Huber

## Abstract

Identifying gene expression differences in heterogeneous tissues across experimental or observational conditions is a fundamental biological task, enabled by single-cell assays such as multi-condition sc-RNA-seq. Current data analysis approaches divide the constituent cells into clusters meant to represent cell types, and identify differentially expressed genes for each cluster. However, such discrete categorization tends to be an unsatisfactory model of the underlying biology. Use of more gradual representations of cell type or cell state promises higher statistical power, better usability and better interpretability. Here, we introduce Latent Embedding Multivariate Regression (LEMUR), a generative model that enables differential expression analysis using a continuous low-dimensional latent space parameterization of cell type and state diversity. It operates without, or before, commitment to discrete categorization. LEMUR (1) integrates data from the different conditions, (2) predicts how each cell’s gene expression would change as a function of the conditions and its position in latent space, and (3) for each gene, identifies compact neighborhoods of cells with consistent differential expression. Unlike statically defined clusters, these neighborhoods adapt to the underlying gene expression changes. We assess LEMUR’s performance on a compendium of single-cell datasets and show applications to the identification of tumor subpopulations with distinct drug responses, the interplay between cell state and developmental time in zebrafish embryos, and the discovery of cell state *×* environment interactions in a spatial single-cell study of plaques in Alzheimer’s disease. LEMUR is broadly applicable as a first-line analysis approach to multi-condition sc-RNA-seq data.

**Software availability:** https://bioconductor.org/packages/lemur

---

Premature discretization of continuous variables leads to artifacts in data analysis and a loss of power; yet, it is the dominant approach to deal with the cell state diversity in multi-condition single-cell data. Lähnemann et al. (2020) described overcoming the reliance on clustering or cell type assignment before downstream analysis as one of the grand challenges in single-cell data analysis. Single-cell RNA-seq can be used to study the effect of experimental interventions or observational conditions on a heterogeneous set of cells, e.g., from tissue biopsies or organoids. Typically, cells from the same sample share the same condition but come from multiple cell types and states (e.g., position in a differentiation or cellular aging path, cell cycle, metabolism). Compared to bulk-sequencing, the novelty of multicondition single-cell RNA-seq is the ability to disentangle expression changes between corresponding cells (i.e., same cell type and state) under different conditions, from those between cell types or states.

Here, we present a generative model and inference procedure to address three tasks in multicondition single-cell data analysis: (1) integrate the data into a common latent space, (2) for each cell, predict the expression it would have in any of the conditions, and (3) find interesting and statistically significant patterns of differential expression. We call our method Latent Embedding MUltivariate Regression (LEMUR).

Many approaches exist for Task 1, with the more or less explicit aim that the variation remaining in the common latent space represents cell type or cell state, and no longer the external conditions. The feasibility of Task 2 depends on whether the function that maps the data to the common embedding space is invertible. This is the case, for instance, for scVI, scGEN, CPA, and CellOT, which are based on parametric autoencoder models (Lopez et al., 2018; Lotfollahi et al., 2019, 2023; Bunne et al., 2023). Other integration methods, including Harmony, Seurat, and MNN (Korsunsky et al., 2019; Haghverdi et al., 2018; Butler et al., 2018), operate nonparametrically, by enumerating the mapped coordinates for all observed cells, and thus have no canonical way to ask the counterfactual.

For differential expression analysis across conditions, the state of the art is to take an integrated embedding, assign the cells to clusters, and find differentially expressed genes separately for each given cluster using methods known from bulk RNA-seq analysis (“pseudobulking”) (Crowell et al., 2020; Missarova et al., 2024). Here, we turn this process around. We address Task 3 by employing the LEMUR counterfactual predictions to compute differential expression statistics for each cell and each gene, and *then* select connected sets of cells with consistent differential expression.

To this end, the LEMUR model decomposes the variation in the data into four sources:

1. the conditions, which are explicitly known,
2. cell types or states that are not explicitly known, but assumed to be representable by a low-dimensional manifold,
3. interactions between the two, and
4. unexplained residual variability.

LEMUR is implemented in an R package on Bioconductor called *lemur* and a Python package called *pyLemur*.

## Results

Figure 1A outlines the LEMUR workflow. The method takes as input a data matrix of size genes *×* cells. It assumes that appropriate preprocessing, including size factor normalization and variance stabilization was performed (Ahlmann-Eltze and Huber, 2023). In addition, it expects two tables of metadata for the cells: a vector which for each cell specifies the sample (independent experimental unit, e.g., tissue biopsy or organoid) it originates from, and a design matrix (Law et al., 2020).

**Figure 1.**
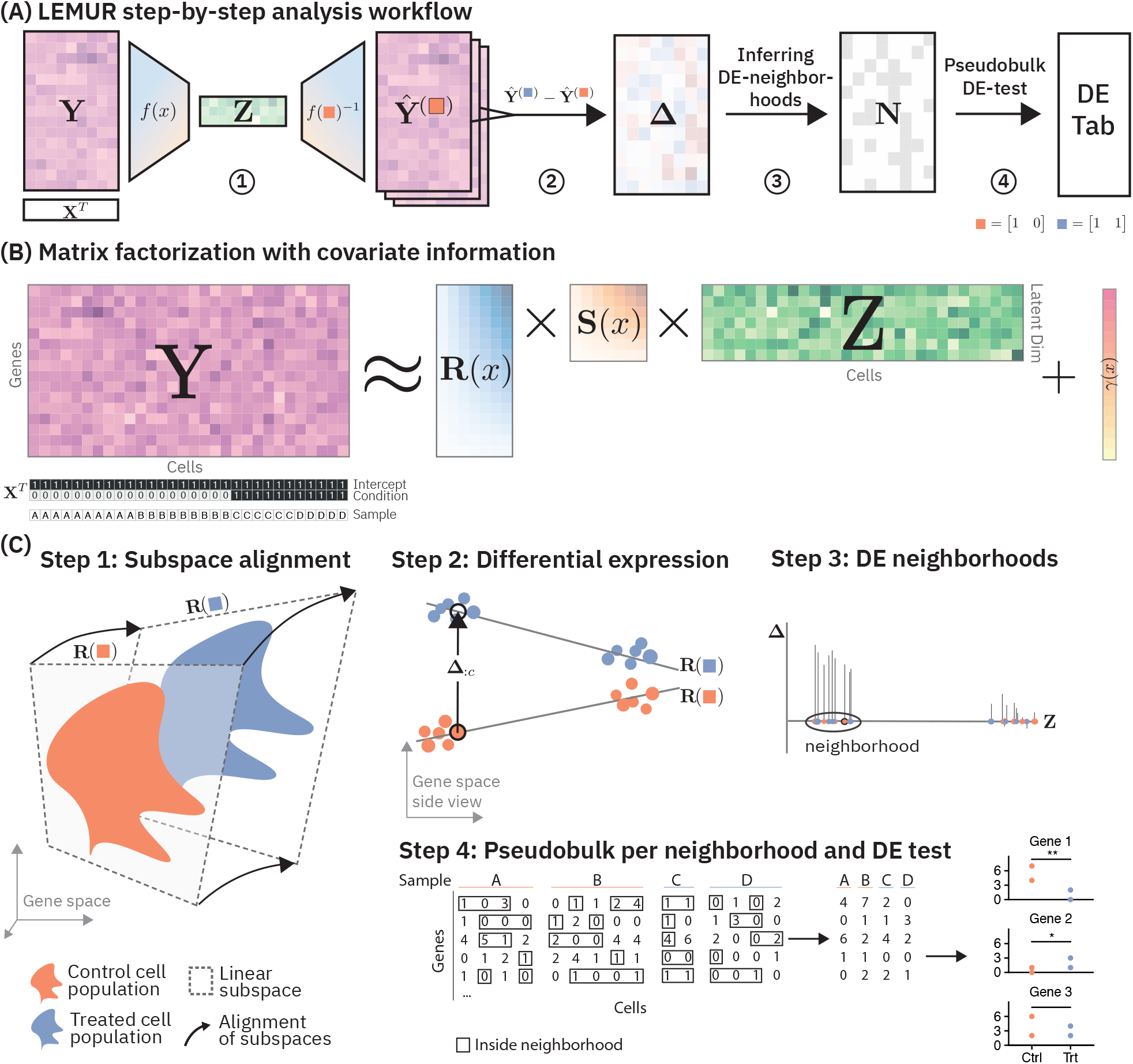
Conceptual overview of latent embedding multivariate regression (LEMUR). (A) Four-step workflow. (B) The matrix factorization at the core of LEMUR. (C) Details on each step from A: Step 1: a linear subspace is fitted separately for each condition. The subspaces for the different conditions are related to each other via affine transformations that are parameterized by the covariates. For this visualization, different 2d subspaces of a 3d gene space are drawn; actual dimensions are higher. Step 2: the differential expression statistic Δ is computed as the difference between the predicted values in the control and treated condition. The visualization shows a top view of the visualization from Step 1. Step 3: for each gene, cells close to each other with consistent Δ values are grouped into neighborhoods. Step 4: a pseudobulk differential expression test is applied to the cells within each neighborhood.

The design matrix encodes one or more covariates that represent experimental treatments or observational conditions. It is analogous to the design matrix in differential expression tools like limma and DESeq2 (Smyth, 2004; Love et al., 2014) and can account for fully general experimental or study designs. The design matrix can include sources of unwanted variation (e.g., experimental batch) and sources of variation whose influence we are interested in (e.g., treatment status). We term a unique combination of covariate values a *condition*. In the simple case of a two-condition comparison, the design matrix is a two-column matrix, whose elements are all 1 in the first column (intercept) and 0 or 1 in the second column, indicating for each cell which condition it is from.

### Matrix factorization in the presence of known covariate information

The central idea of LEMUR is a multi-condition extension of principal component analysis (PCA) (Fig. 1B). Given a data matrix **Y**, PCA can be used to approximate **Y** ≈ **RZ** with two smaller matrices: the first, **R**, is a basis that spans a suitable low-dimensional linear subspace of the original data space, the second, **Z**, contains the coordinates of each cell with respect to that basis. In this elementary form, there is no place to explicitly encode known experimental or study covariates. LEMUR adds this capability by including a regression analysis component.

Instead of using a single subspace, we let the subspace spanning matrix **R**(**X**) depend on the covariates provided in the design matrix **X** (Fig. 1C, step 1). With this ansatz we address the decomposition task posed in the introduction: known sources of variation are encoded in **X**, cell type and state variations are represented by the cell coordinates **Z**. While **X** is explicitly known, **Z** is latent, i.e., is estimated from the data. The construction allows modeling interactions, that is, gene expression changes across conditions that are different for different cell types and states. This is the main feature of LEMUR.

A second feature of this construction is between-condition integration: data from cells observed in different conditions are mapped into a common latent space. By default, the integration is based on the alignment of the respective subspaces, but it can optionally be improved by information that indicates that certain cells observed in different conditions correspond to each other and thus should be close to each other in the latent space **Z**. Such information can come in the form of explicit landmarks, i.e., from cells that express distinctive marker genes, or via statistical properties of the cells exploited by methods such as Harmony (Korsunsky et al., 2019).

We account for such correspondence by adding an affine transformation **S**(**X**) to the model.

A third feature of our model is its ability to predict, for each cell, how its gene expression profile would look like in any of the conditions— even though it was only observed in one of them. In fact, predictions are available not only for those positions in latent space where cells were observed, but for all positions, i.e., also for hypothetical interpolating or extrapolating cell types and states. We use these predictions to find changes of gene expression that are coordinated across regions of latent space, i.e., across the same or similar cell types and states.

Fig. 2 shows a stylized illustration of the LEMUR approach. LEMUR fits one 1D subspace (line) per condition, each parameterized by a rotation applied to a common base space. The parametric model yields a predicted expression value for each cell in each condition, and we look for regions in latent space—here, we have two major regions, left and right—in which predictions are consistently positive or negative. In the Methods section, we provide a more formal mathematical specification.

**Figure 2.**
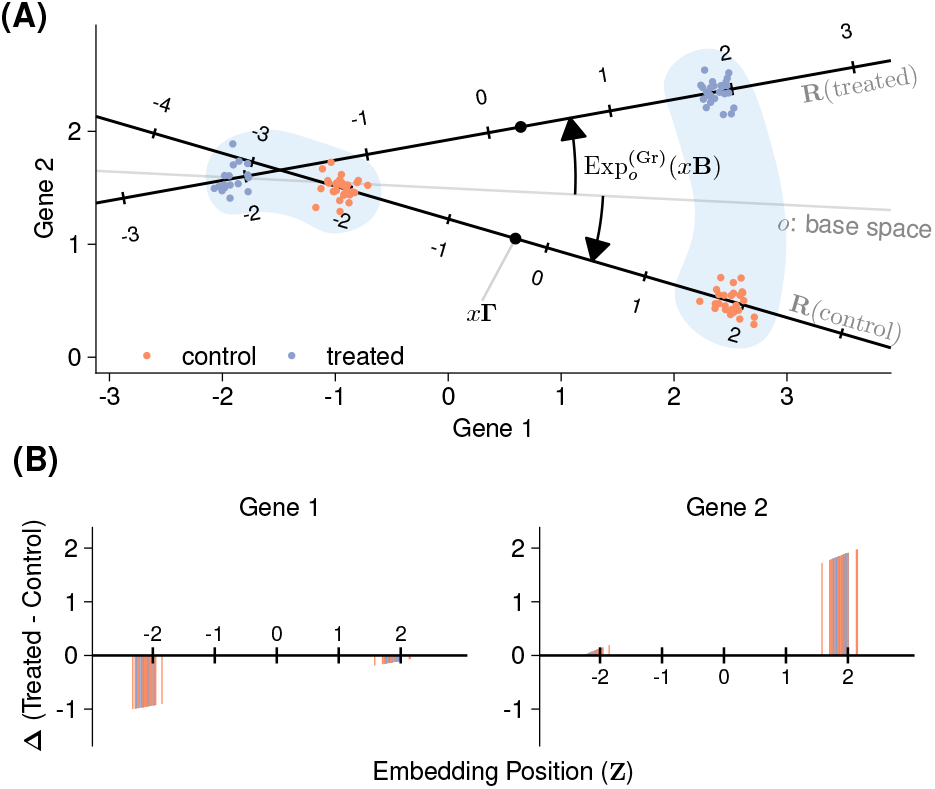
Stylized example with two genes, observed in two groups of cells, in two conditions. (A) Scatter plot of the genespace with condition-specific one-dimensional subspaces. See the Methods for the mathematical details. (B) Predicted expression of the two genes in each cell. Gene 1 is lower expressed in treatment, compared to control, only in the “left” cell type, Gene 2 is up in the treatment, only in the “right” cell type.

### Cluster-free differential expression analysis

We can predict the expression of a cell in any condition using this parametric model. The differential expression between two conditions (Fig. 1C, step 2) is just the difference between their predictions and can be computed for each cell—even though for any cell, data was only observed in exactly one replicate of one condition (Fig. 2B). The resulting matrix of differential expression estimates **Δ** has two uses: first, we can visualize the differential expression values for each gene as a function of latent space. Typical choices for the dimension of the latent space are ten to a hundred and for its visualization we use a further non-linear dimension reduction into 2D scatterplots, such as UMAP (McInnes et al., 2018). Examples are shown in Figs. 4 and 6. Second, we use **Δ** to guide the identification of differential expression neighborhoods, i.e., groups of cells that consistently show differential expression for a particular gene (Fig. 1C, Step 3). The intention for these neighborhoods is to be connected, convex, and maximal, i.e., the differential expression pattern would become disproportionately less consistent and less significant if the neighborhood were extended.

To rank and assess our level of confidence in the found neighborhoods, we do not attempt to measure the statistical uncertainty of the predictions **Δ**. Instead, we use pseudobulk aggregation (Crowell et al., 2020) of the original data: raw counts, if available, otherwise the log-normalized values. For each sample, we sum up the original counts or take the mean of the log-normalized values of the cells in the neighborhood to obtain a neighborhood-specific genes *×* samples table (Fig. 1C, Step 4), followed by a differential expression test with glmGamPoi, edgeR, or limma (Ahlmann-Eltze and Huber, 2020; Robinson et al., 2010; Smyth, 2004).

### Outputs

LEMUR produces the following outputs:

- a common low-dimensional latent space representation of all cells (**Z**),
- parametric transformations **R** and **S** that map the condition-specific latent spaces into each other,
- the predicted expression **Ŷ** and cell in any condition, for each gene
- the predicted differential expression **Δ** for each gene and cell, for any contrast that can be constructed from the design matrix,
- for each gene and contrast, a neighborhood of cells and a statistical measure of significance (p value).

The last of these will usually be the one end users care about most; the others are useful for diagnostics, quality assessment, visualization, and further modeling uses of the data.

The explicit parameterization of the transformations **R** and **S** means that they can easily be inter- or extrapolated beyond the observed set of data, and inverted from the low-dimensional embedding back to the data space. In this sense, LEMUR is a generative model.

### Performance assessment

We assessed performance of LEMUR using 13 publicly available multi-condition single-cell datasets (listed in Availability). We preprocessed each dataset consistently following Ahlmann-Eltze and Huber (2023).

First, we considered integration performance: how well does the joint low-dimensional representation of the cells preserve biological signal encoded in the latent space, but remove traces of the known covariates such as batch and treatment effects. Figure 3A illustrates this on the dataset by Kang et al. (2018) of Lupus patient samples treated with either interferon-*β* or vehicle control. We measured the covariate removal by counting for each cell how many of its *k* = 20 neighbors come from the same condition (*k*-nearest neighbor (*k*-NN) mixing). For a balanced dataset with two conditions, an ideal method scores a *k*-NN mixing value of *k/*2 = 10. We measured the biological signal retention by comparing—for each condition separately—a clustering of the embedding with a clustering of the original data, as measured by the mean of the two adjusted Rand indexes (ARI). An ideal method scores close to ARI = 1.

**Figure 3.**
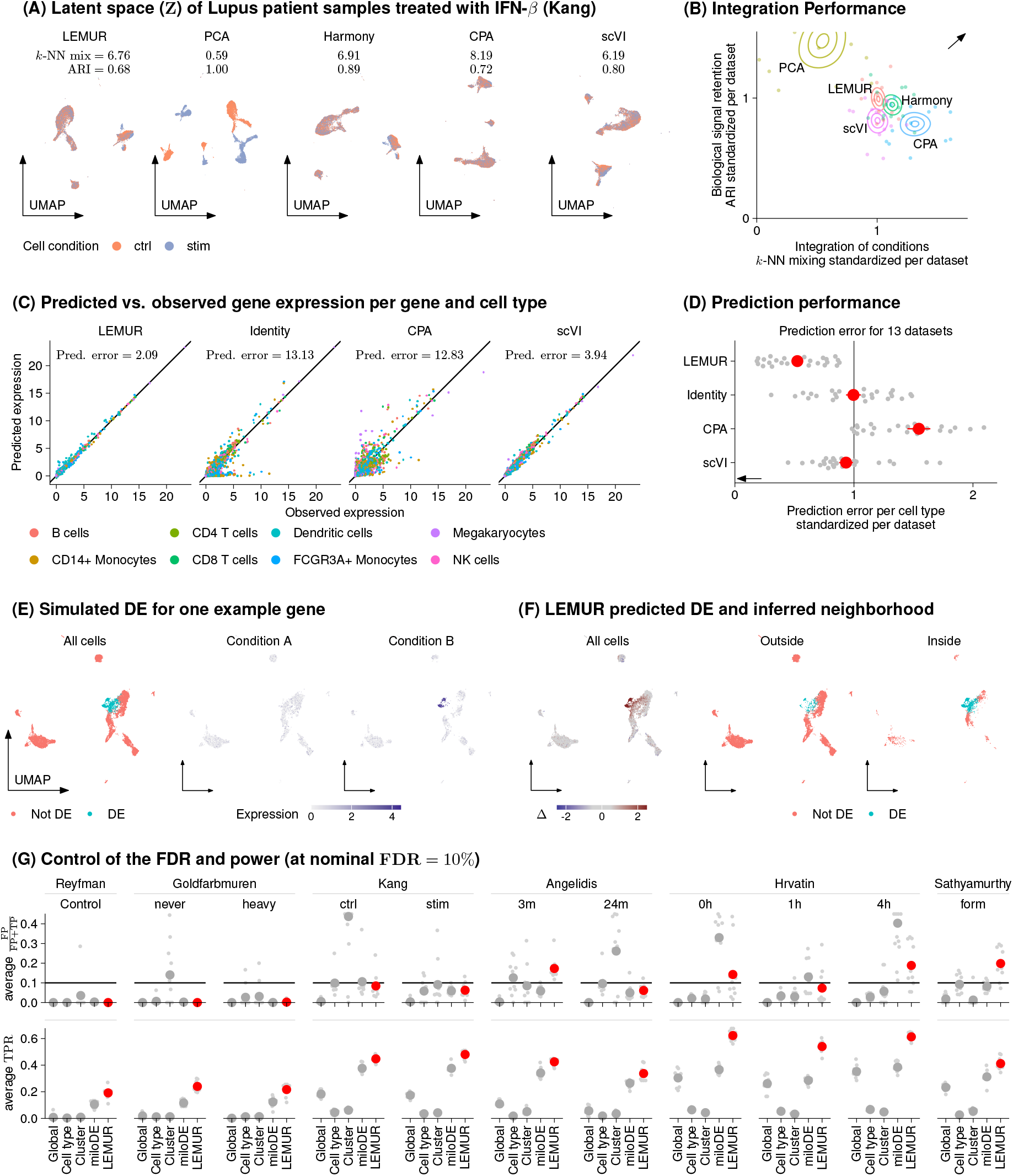
Performance assessment. (A) UMAPs of the latent spaces for the data by Kang et al. (2018). The *k*-NN mixing coefficient and the ARI are defined in the main text. (B) Density plots of the bootstrapped mean performance for ARI and *k*-NN mixing across thirteen datasets. To adjust for the dataset-dependent variation, we divided the *k*-NN and ARI score by the average per dataset. (C) Scatter plots of predicted expression under treatment (*y*-axis) against observed expression (*x*-axis), for 500 genes in each of 8 cell types (same data as in A). (D) Prediction error as in C, across the same thirteen datasets as in B (grey points). Red points show the mean. (E) Simulation setup. For one of the implanted genes, the left panel shows a UMAP of the LEMUR latent space, where the color indicates whether an expression change in this gene was simulated for that cell. The center and right panels show the simulated expression values. (F) Left panel: predicted log fold change for the gene from panel E (**Δ** = Ŷ^B^ − Ŷ^A^). Center and right panels: the set of cells inside or outside the inferred neighborhood. (G) Comparison of observed false discovery proportion and true positive rate (TPR) for 11 datasets, with 10 replicates and the overall mean shown as a large point. The nominal FDR was fixed to 10.

Across the 13 datasets, the performance of LEMUR on these measures was similar to that of Harmony (Fig. 3B). Other methods make different trade-offs between the two measures, and no method clearly dominates. Suppl. Fig. S1 shows that the results are consistent across 7 additional metrics.

The computational cost of running LEMUR is at the low end of what may be expected for such data, and comparable to that of other approximative PCA methods. For instance, computing the first 50 latent dimensions on the Goldfarbmuren data, with 24 178 cells and 20 953 genes (which occupies 4 GB of RAM), took us 35 seconds and 24 GB RAM with the approximative IRLBA algorithm (Baglama et al., 2022). For comparison, fitting LEMUR without integration took 103 seconds and needed 33GB RAM. Aligning the cells with landmarks or Harmony added 2 and 95 seconds, respectively (Suppl. Fig. S2).

Next, we assessed the ability of LEMUR to predict gene expression across conditions. We used it to predict gene expression under treatment for cells that were observed in the control condition, and compared these predictions to the data from cells that, in fact, were treated. To avoid overfitting, we assessed predictions on “hold-out” cells whose data were not used for training. As there is no direct correspondence between individual cells observed under the two conditions, we considered averages across annotated cell types. Fig. 3C shows scatter plots of the predicted–observed comparison for the Kang et al. (2018) dataset for the 500 most variable genes in eight cell types for four methods: CPA, scVI, LEMUR, and the trivial prediction of no change (Identity). Across the thirteen datasets, LEMUR showed the smallest prediction error measured by the *L*_2_ distance between observed and predicted values (Fig. 3D). Suppl. S3 shows that the results are consistent across six additional metrics.

In a third set of comparisons, we tested the ability of LEMUR to identify sets of cells with consistent differential expression. We took all cells from the control condition of the Kang et al. (2018) data, assigned them randomly to a condition *A* or *B*, and implanted genes with differential expression in a subset of cells. Fig. 3E shows an example. LEMUR accurately identified the expression change and inferred a neighborhood of cells that overlapped well with the simulated ground truth (Fig. 3F).

We expanded this analysis to ten more semisynthetic datasets, each with 200 implanted differentially expressed genes, to asses the type I error control of the LEMUR differential expression test. LEMUR on average controlled the false discovery rate (FDR) (Fig. 3G upper panel). In addition, it was more powerful than a pseudobulked test across all cells (*global*), or separate tests for subsets of cells, either by *cell type* or *cluster* as in Crowell et al. (2020), or by neighborhood as in *miloDE* (Missarova et al., 2024) (Fig. 3G, lower panel). Suppl. Fig. S4 assesses FDR control and power for additional variants of the LEMUR method. It shows that accounting for the post-selection bias from the neighborhood inference is important, and that it is successfully addressed by the data splitting.

Overall, these benchmarks demonstrate that LEMUR (1) successfully integrates single-cell data from different conditions, (2) detects cell type/state specific differential expression patterns without access to prior clusterings or categorizations of cells, and (3) provides accurate statistical type I error control and good power. In the following, we apply LEMUR to the analysis of different biological datasets.

### Analysis of a treatment/control experiment: panobinostat in glioblastoma

Zhao et al. (2021) reported single-cell RNA-seq data of glioblastoma biopsies. Aliquots from five patients were assayed in two conditions: control and panobinostat, a histone deacetylase (HDAC) inhibitor (Fig. 4A). After quality control, the data contained 65 955 cells (Suppl. Tab. S1).

**Figure 4.**
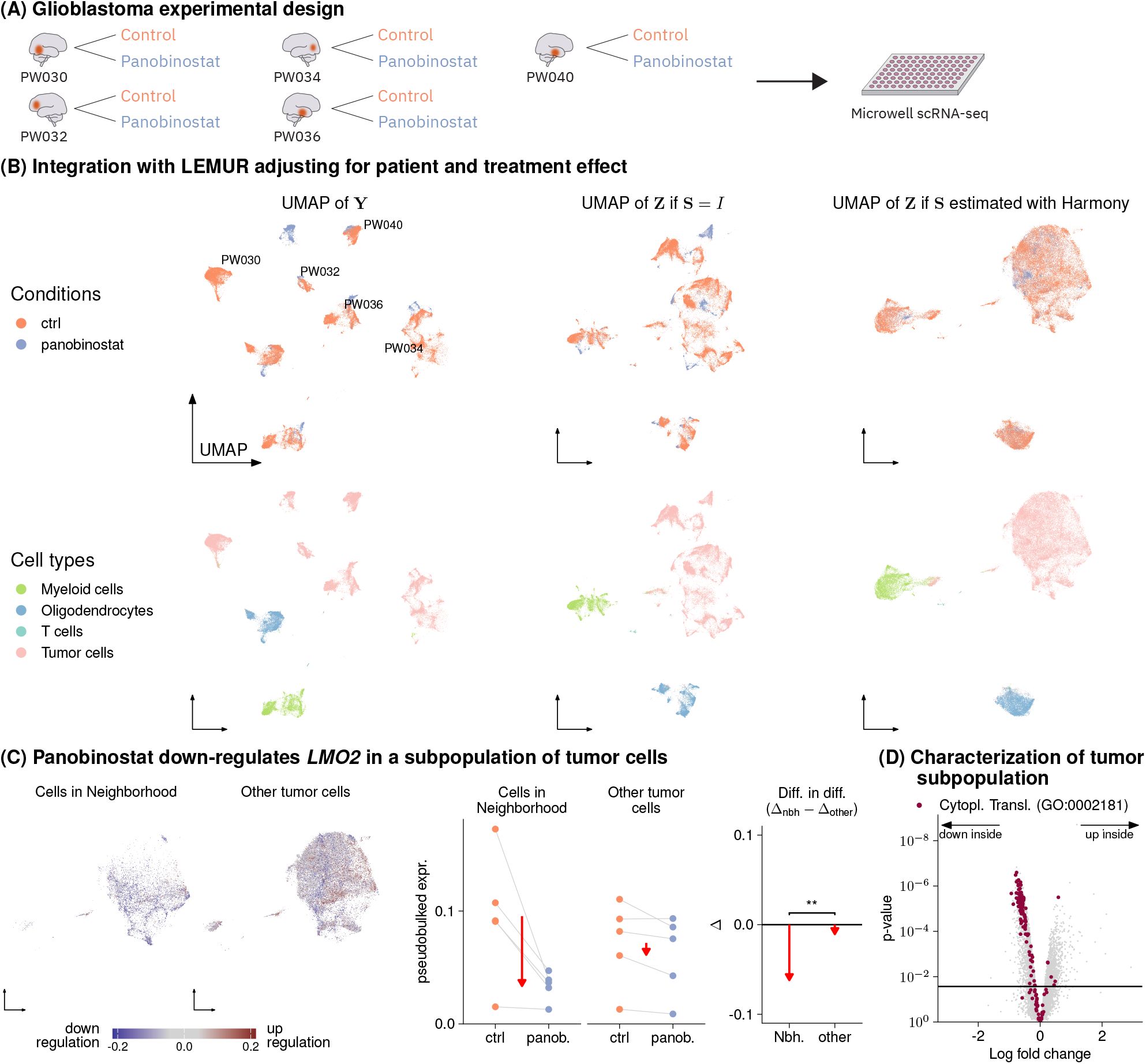
Analysis of five glioblastoma tumor biopsies. (A) Experimental design. (B) UMAP of the log-transformed data (first column) and the latent embeddings produced by LEMUR (second and third column), colored by treatment and cell type (first and second row, respectively). (C) Differential expression analysis of *LMO2* within the tumor cells. Faceted UMAP of the cells inside and outside the neighborhood. The scatter plots in the middle show the pseudobulked expression values from cells inside and outside the neighborhood by donor and condition. The panel on the right compares the differences inside and outside the neighborhood (red arrows). (D) Volcano plot for the comparison between cells in- and outside the subpopulation from panel F, all in the control condition. Gene set enrichment analysis points to reduced translation activity in the subpopulation.

The left column of Fig. 4B shows a twodimensional UMAP of the input data **Y**. Most visible variation is associated with the known covariates: donor and treatment condition. Some further variation is related to the different cell types in the biopsies. We used LEMUR to absorb donor and treatment effects into **R**, setting the latent space dimension to *P* = 60. The middle column of Fig. 4B shows, upon fixing **S**(*x*) = *I*, a UMAP of the resulting matrix **Z** of latent coordinates for each cell. Cells from different samples are more intermixed, and within-sample cellular heterogeneity is more evident. This picture becomes even clearer after using **S** to match cell subpopulations across samples using Harmony’s maximum diversity clustering (Fig. 4B, right column). Here, a large population of tumor cells and three non-tumor subpopulations become apparent.

The successive improvement of the latent space representation from left to right is further demonstrated in the lower row of Fig. 4B, where the points are colored according to a cell type assignment that we obtained from the expression of selected marker genes and known chromosomal aneuploidies (Suppl. Fig. S5A,B).

The linear latent space of LEMUR is readily interpretable. This is exemplified in Suppl. Figure S5C, which extends the biplot concept from PCA (Gabriel, 1971) to the multi-condition setting. We can explore how higher or lower expression of any gene affects a cell’s position in the latent space **Z** by plotting the gene’s loading vector relative to the coordinate system of **Z**.

Using LEMUR’s differential expression testing, we found that panobinostat caused cell subset-specific expression changes in 25% of all genes (2 498 of 10 000) at an FDR of 10%. Suppl. Fig. S6 shows the differential expression and inferred neighborhoods for seven genes.

Focusing on the tumor cells, we identified subpopulations that differentially responded to panobinostat treatment (Fig. 4C). In a subpopulation of 9 430 tumor cells, which stemmed in significant proportions from all five patients, treatment with panobinostat caused downregulation of *LMO2*, while in the majority of tumor cells (*n* = 36 535) expression of *LMO2* was unchanged (Fig. 4C, right panel). *LMO2* forms protein complexes with the transcription factors *TAL1, TCF3*, and *GATA*; it is important for angiogenesis and was originally discovered as an oncogene in T cell acute lymphoblastic leukemia (TALL) (Chambers and Rabbitts, 2015). Kim et al. (2015) studied the role of *LMO2* in glioblastoma and found that higher expression is associated with worse patient survival. They concluded that *LMO2* could be a clinically relevant drug target. To further characterize the subset of tumor cells that respond to panobinostat by downregulating *LMO2*, we compared their overall gene expression profiles in the control condition to that of the other tumor cells. We found lower abundances of ribosomal genes, consistent with lower translational activity (Fig. 4D).

### Analysis of a time course: zebrafish embryo development

Saunders et al. (2023) reported an atlas of zebrafish embryo development, which includes data from 967 embryos and 838 036 cells collected at 16 time points in 2h intervals from 18h to 48h post fertilization (Fig. 5A). They used their scRNA-seq data to assign each cell to a common cell type classification scheme and studied the temporal dynamics of appearance and disappearance of cell types along developmental time. We asked whether gene expression changes could reveal additional biological phenomena. Thus, we looked for temporal profiles of gene expression that systematically differed across cells that shared the same cell type annotation. For this, we used the ability of LEMUR to predict any gene’s expression at any point in latent space at any time. To represent the time dependence of each gene with a smaller number of parameters, we used natural cubic splines with three degrees of freedom, following Smyth et al. (2023). To model latent space variation, we interpolated linearly in latent space **Z** between two cells (red and green) from the central nervous system and analogously, between two cells (purple and turquoise) from the periderm, a transient, outermost epithelial layer that covers the developing embryo during the early stages of development (Fig. 5B, Suppl. Fig. S7A).

**Figure 5.**
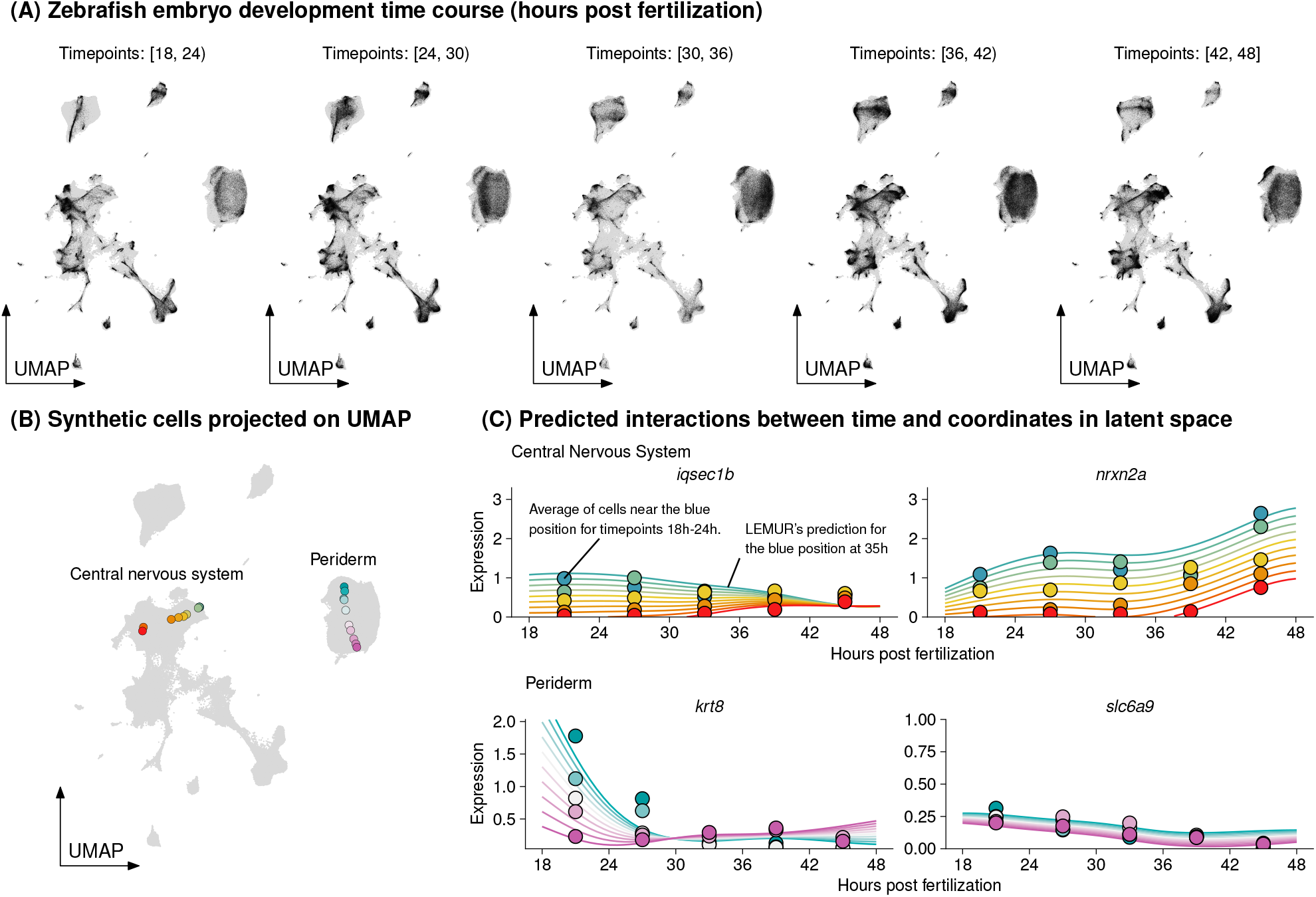
Analysis of a time course single-cell experiment. (A) UMAPs of embryonic development based on the integrated latent space of LEMUR. Black: cells from the respective time window post fertilization; grey: all other cells, for comparison. Since some cell types only exist at particular stages of development, temporal changes in the distribution of black points are to be expected. (B) Synthetic cells projected onto the UMAP. They interpolate between pairs of observed cells, one pair in the periderm (purple/turquoise) and one in the central nervous system (red/green). (C) Expression predictions (smooth lines) and averaged observed data (points) for four genes as a function of time (*x*-axis) and latent space coordinates (color).

We used the LEMUR fits to screen for genes for which the spline coefficients were different across this interpolated latent space gradient, according to a statistical test for interaction (FDR=0.001). Fig. 5C shows the data for four examples in which the temporal divergence of gene expression was corroborated by pseudobulking the observed expression data of the nearest neighbors in each 6hour interval. For instance, *krt8*, which encodes a keratin essential for the structural integrity, protective barrier function and proper development of the periderm, showed decreasing expression over time for cells close to the turquoise cell, and increasing expression for cells close to the purple cell. Several possible explanations for the intricate and divergent temporal regulation of *krt8* within the periderm cells (Suppl. Fig. S7B,C) exist, including spatial structures.

### Analysis of spatially-resolved expression: plaque density in Alzheimer’s disease

Cable et al. (2022) performed Slide-seq V2 on the hippocampus of four mice genetically engineered to model amyloidosis in Alzheimer’s disease. Using microscopy, they quantified the spatial density of Amyloid *β* plaques (Fig. 6A). Thus, plaque density is an observational covariate that varies from cell to cell; this is in contrast to covariates considered above, which vary from sample to sample. Cable et al. (2022) reported a differential expression analysis per discrete cell type category; however, the categories were fairly broad and could not account for the gradual changes suggested by the data (Fig. 6B, Suppl. Fig. S7A). LEMUR enabled us to defer cell type categorization and directly identify genes whose expression varied between low and high plaque density in adaptively found subsets of cells. Fig. 6C shows the differential expression prediction for *Jun*, a transcription factor that was identified as a member of the pathway regulating *β* amyloid-induced apoptosis (Morishima et al., 2001; Akhter et al., 2015) and one of the top hits after the LEMUR analysis. The correlation between higher Amyloid-*β* plaque density and increased *Jun* expression was limited to a subset of about 20% of the neurons, which clustered both in spatial coordinates and in the latent space (Fig. 6D). The correlation did not hold for other neurons (Fig. 6E, Suppl. Fig. S8B). We projected the data from Cable et al. onto the hippocampus reference atlas by Yao et al. (2021) and found that this subset belonged to the glutamatergic neurons from the dentate gyrus and CA1 (prosubiculum) (Suppl. Fig. S8A).

**Figure 6.**
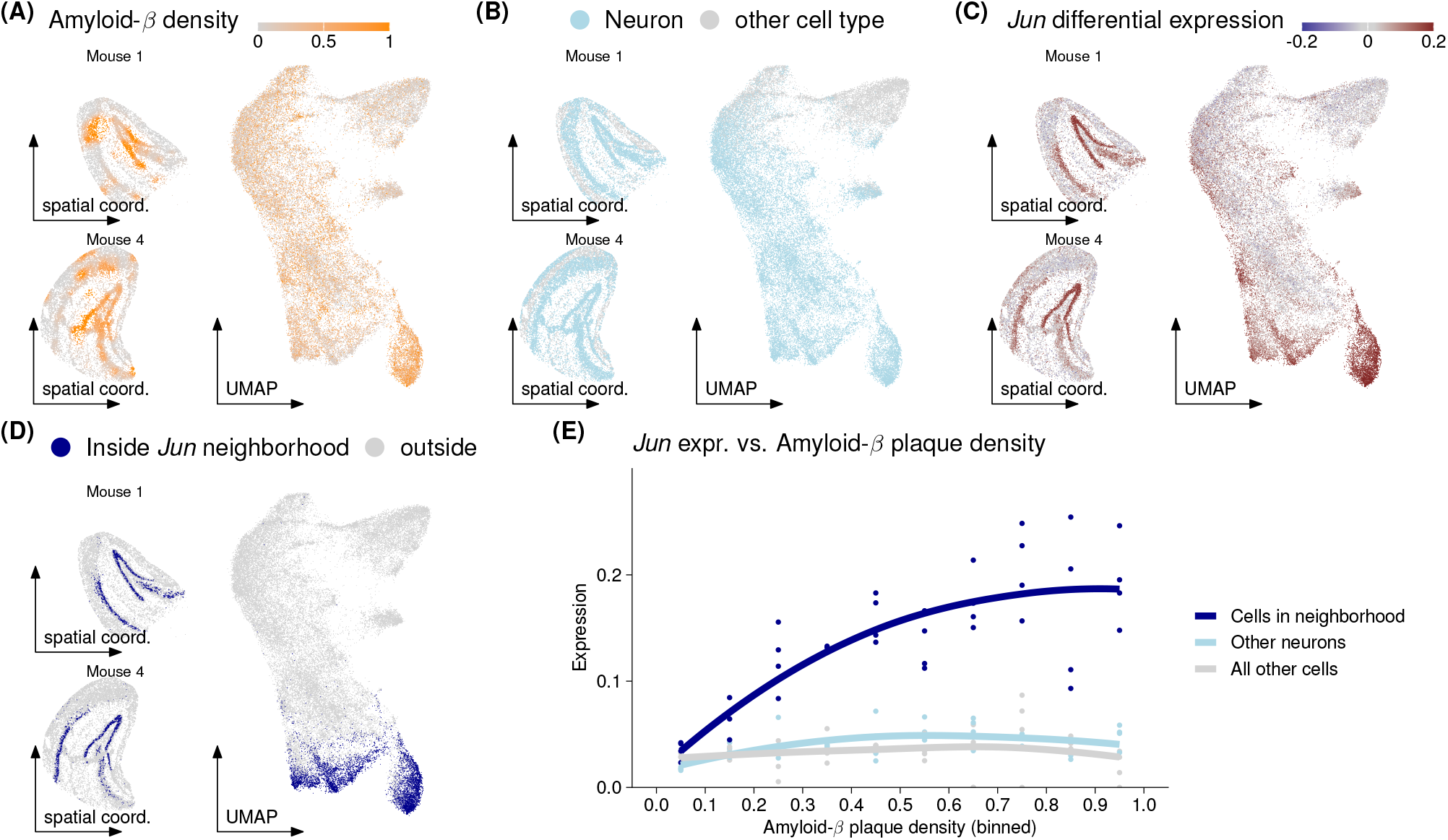
Analysis of a spatial single-cell experiment. (A-D) UMAPs of the cells from the hippocampi of four mice, and spatial maps for two of them, colored by (A) Amyloid-*β* plaque density, whether the cell is a neuron, (C) the differential expression for *Jun*, and (D) whether the cell is inside the differential expression neighborhood for *Jun*. (E) Scatter plot of *Jun* expression as a function of plaque density for cells inside the neighborhood, neurons not in the neighborhood, and all other cells. The line is a LOESS fit through the pseudobulked values.

## Discussion

Latent embedding multivariate regression (LEMUR) enables differential expression analysis of single-cell resolution expression data between composite samples, such as tissue biopsies, organs, organoids or whole organisms. The method allows for arbitrary experimental or study designs, specified by a design matrix, just as in ordinary linear regression, or in ‘omics-oriented regression methods like limma (Smyth, 2004), edgeR (Robinson et al., 2010), DESeq2 (Love et al., 2014). Applications range from comparisons between two conditions with replicates, over paired studies, such as a series of tissue biopsies before and after treatment, over studies with multiple covariates (e.g., genetic and drug perturbations), interactions between covariates, and continuous covariates. The method represents cell-to-cell variation within a condition using a continuous latent space. Thus, it avoids—or postpones—the need for categorical assignment of cells to discrete cell types or cell states, and offers a solution to one of the challenges identified by Lähnemann et al. (2020).

We demonstrated the utility of the approach on three prototypical applications, and we benchmarked important aspects of its performance on a compendium of 13 datasets. The application cases are a matched control-treatment study of patient samples in glioblastoma, an atlas of Zebrafish embryo development where time is a continuous covariate, and a spatial transcriptomics study of Alzheimer’s plaques where plaque density is a continuous covariate. We showed how LEMUR identified biologically relevant cell subpopulations and gene expression patterns.

To achieve this, we combine latent space representation by dimension reduction with regression analysis in a novel matrix factorization approach. The model is predictive: for each observed cell, it predicts its gene expression in any of the conditions, even though it was only measured exactly once. Moreover, as each cell is parameterized by a position in *P* -dimensional real vector space, the model can also predict the expression of “synthetic cells” at unobserved positions, say in-between observed cells or extrapolating out, in any condition. We use these capabilities for differential expression analysis.

We detect neighborhoods of cells with consistent differential expression patterns with respect to comparisons (“contrasts”) of interest. The neighborhoods are found in a data-driven manner. No a priori categorization of cells into “cell types” is needed, but once neighborhoods have been identified, one can annotate or compare them with whatever annotation that is relevant.

Our current implementation of neighborhood finding leaves room for future improvements. It is stochastic, by relying on a random sample of one-dimensional projections of point clouds in *P* -dimensional space. Thus, repeated running of the algorithm can result in (slightly) different outputs. Also, it addresses the post-selection inference problem using a rather heavy-handed data splitting approach.

Unlike some other single-cell data integration and expression prediction tools, LEMUR is built around linear methods. It is parameterized with a modest number of parameters and uses a small number of layers. This is in contrast to the often-repeated claim that the complicatedness of single-cell data necessitates non-linear methods and ‘deep’ models. We showed that our approach based on simple, linear matrix decomposition using a sufficiently high-dimensional latent space is capable of representing the data in a useful manner. Compared to deep-learning approaches, LEMUR’s interpretable and easy-to-inspect model should facilitate dissection and follow-up investigation of its discoveries. In a Supplementary Note, we discuss how the LEMUR model differs from other approaches that combine dimension reduction and regression.

Overall, we believe that LEMUR is a valuable tool for first-line analysis of multi-condition single-cell data. Compared to approaches that require discretization into clusters or groups before differential expression analysis, representing cell types and states in a continuous latent space may better fit the underlying biology, and may facilitate the precise identification of affected cells. This in turn should ease analysts’ work, and enable biological discoveries that could otherwise be missed.

## Availability

All datasets used in this manuscript are publicly available:

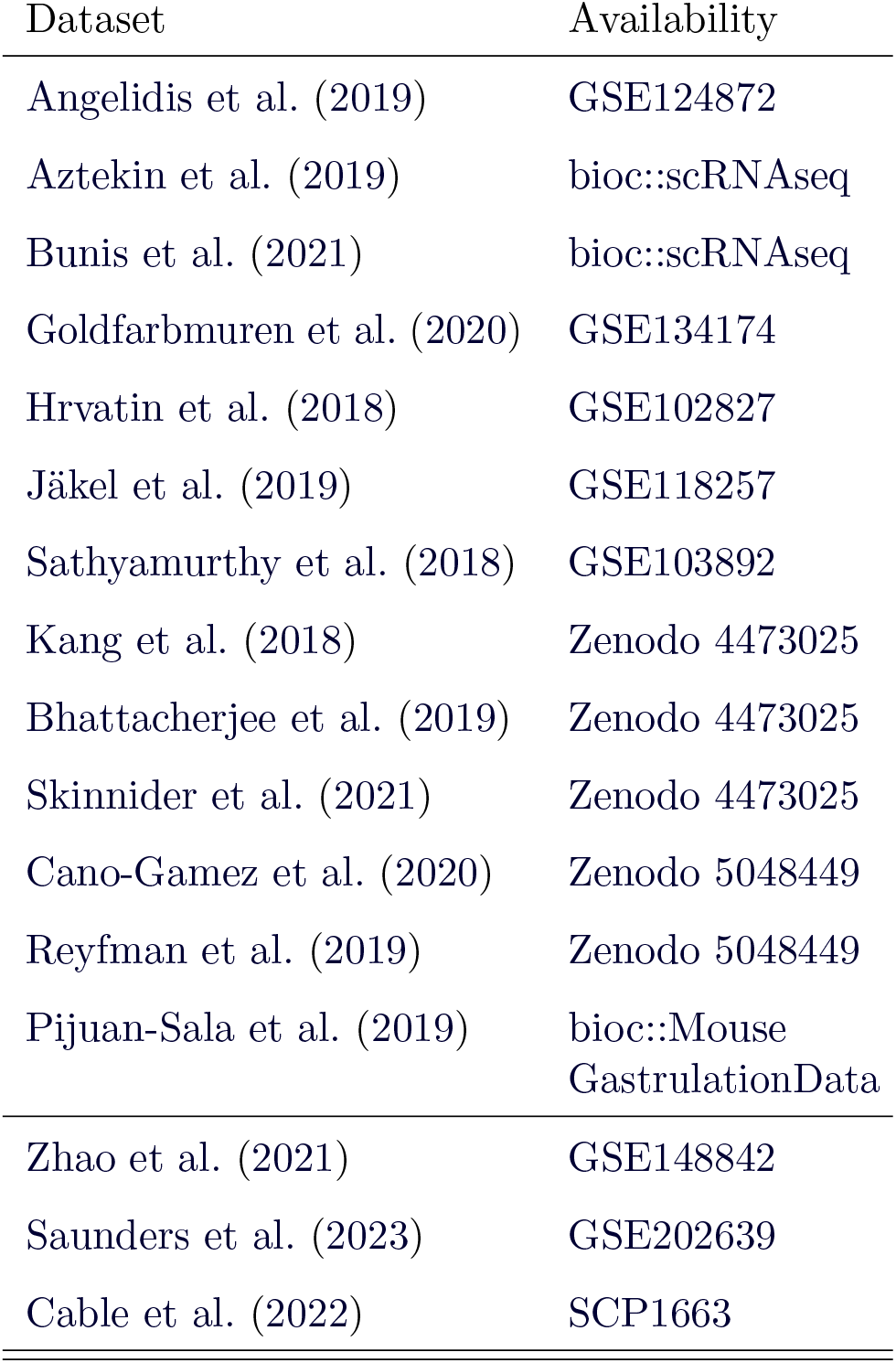

The *lemur* R package is available at bioconductor.org/packages/lemur and the code to reproduce the analysis is available at github.com/const-ae/lemur-Paper which we also permanently archived using Zenodo doi.org/10.5281/zenodo.12726369. A Python implementation of the LEMUR model (without the differential testing capabilities) is available at github.com/const-ae/pylemur.

## Acknowledgments

We thank Simon Anders, Oliver Stegle and Judith Zaugg for valuable feedback and discussions. We thank Ronny Bergmann for his advice on optimization on manifolds. We thank Lauren Saunders for early access to and discussion of the zebrafish data. We thank EMBL IT services and HPC resources for providing infrastructure to perform computations needed for this work.

We thank EMBL Kinderhaus for providing authors with thinking time through their excellent childcare.

## Methods

Given the data matrix **Y** of size *G × C*, where *G* is the number of genes and *C* is the number of cells, consider the decomposition

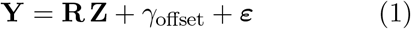

with the *G × P* -dimensional matrix **R**, the *P × C* matrix **Z**, the *G × C* matrix ***ε***, the *G*dimensional vector *γ*_offset_ (all real-valued) ^1^, and *P <* min(*G, C*). To simplify interpretation, we require the columns of **R** to be orthonormal (i.e., they are an orthonormal basis of a *P* –dimensional linear subspace of ℝ^*G*^). The embedding **Z** can then be considered the *P* -dimensional coordinates of *C* points in that linear subspace, each representing a cell. Setting *γ*_offset_ to the rowwise means of **Y**, the matrices **Z** and **R** are fit by minimizing the sum of squared residuals

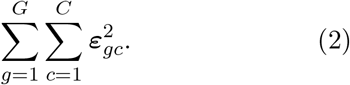

Principal component analysis (PCA) is a special case where, in addition, the columns of **R** are obtained from an eigendecomposition and ordered by eigenvalue. Alternatively, PCA can be understood as the decomposition where **R** is orthonormal and **Z** is orthogonal, which emphasizes the relation to singular value decomposition. In the applications considered in this work, *P ≪* min(*G, C*), and **R** and **Z** can be considered a lower-dimensional approximation of the full data matrix **Y** − *γ*_offset_.

We extend model (1) to incorporate known covariates for each cell. Thus, we consider not just a single matrix **R** and a single vector *γ*_offset_, but treat them as functions of the covariates,

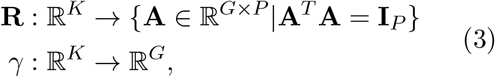

where the arguments of these functions are rows of the *C × K* design matrix **X**, i.e., elements of ℝ^*K*^. The output **R**(*x*) is an orthonormal *G × P* matrix and the output *γ*(*x*) is a ℝ^*G*^ vector. Model (1) then becomes, for each cell *c*,

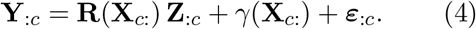

Setting *γ*(*X*_*c*:_) to the least sum of squares solution of regressing **Y** on **X**, the matrix **Z** and **R** ∈ *ℛ* are fit by minimizing the sum of squared residuals (2), where *ℛ* is a suitable set of matrix-valued functions that we define in the following.

Model (4) with these additional features can be considered a multi-condition extension of PCA. Intuitively, this *multi-condition PCA* finds a function **R** that generates for each condition (for each distinct row of the design matrix **X**) a *P* -dimensional subspace that minimizes the distance to the observed data in that condition by rotating a common *base space* into the optimal orientation (Fig. 1C Step 1). **Z** is the orthogonal projection of the data on the corresponding subspace. Stability is ensured by constraining **R** to come from a set *ℛ* of “well-behaved”, smooth functions of the covariates.

To construct *ℛ*, we need to recall some concepts from differential geometry. Given the whole numbers *G* and *P*, the set of all orthonormal real matrices of dimension *G × P* is a differentiable manifold, the Stiefel manifold *V*_*P*_ (ℝ^*G*^). For our application, it is appropriate to consider two matrices equivalent iff they span the same linear subspace of ℝ^*G*^. The set of all such equivalence classes is again a differentiable manifold, called the Grassmann manifold Gr(*G, P*) (Edelman et al., 1998; Bendokat et al., 2023). Accordingly, elements of Gr(*G, P*) are *P* -dimensional linear subspaces of the ambient space ℝ^*G*^. Computationally, we represent an element of Gr(*G, P*) by an orthonormal matrix, i.e., by one of the members of the equivalence class.

We then construct *ℛ* as the set of all functions **R** that have domain and codomain as in Eqn. (3) and can be written as

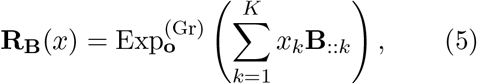

where **B** is a 3-dimensional real-valued tensor of size *G × P × K*. The expression Exp^(Gr)^ is the exponential map on the Grassmann manifold. It takes a point **o** ∈ Gr(*G, P*) and a tangent vector at that point, and returns a new point on the Grassmann manifold. Thus, given a choice of the point **o**, which we call the *base point*, and of the design matrix **X**, the set of all possible **B** induces *ℛ*, and fitting Model (4) is achieved by fitting **B**. We use the terms *tangent vector* and *tangent space* in their standard meaning in differential geometry and represent tangent vectors with *G × P* matrices. The name *exponential map* derives from the fact that for some Riemannian manifolds the exponential map coincides with the matrix exponential; however, this is not the case for Grassmann manifolds. Here the exponential map for a base point **o** and a tangent vector represented by **A** ∈ ℝ^*G×P*^ is

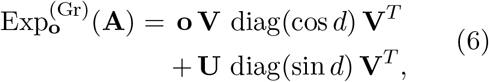

where **A** = **U** diag(*d*) **V**^*T*^ is the singular value decomposition of **A** (Edelman et al., 1998)

The argument of the exponential map in Eqn. (5) is a linear combination of the slices of **B**. Each slice of **B**_::*k*_ represents a tangent vector (**B**_::*k*_ ∈ *𝒯*_**o**_Gr(*G, P*)) and so are their linear combinations (a tangent space is a vector space).

We analogously parameterize the offset function *γ*(*x*) = Σ_*k*_ **Γ**_: *k*_*x*_*k*_, where **Γ** ∈ ℝ^*G×K*^. Accordingly, fitting *γ* is just ordinary linear regression.

### Non-distance preserving extension

Multi-condition PCA (Eqn. (4)) fits subspaces that approximate the data for each condition, but does not depend on shape, scale, or more generally, the statistical distribution of the cells’ embeddings in that subspace. Therefore it can also not adjust for (“absorb”) such differences. This rigidity increases stability and can be a desirable model feature for some applications, by preventing overfitting, but for other applications it can also be a limitation. We extend Model (4) with an optional term **S**, a non-distance preserving, affine isomorphism of ℝ^*P*^, to (i) obtain additional flexibility and (ii) enable input of prior knowledge and user preferences in cell matching:

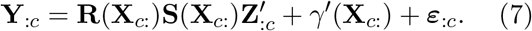

Here, 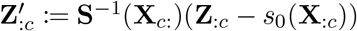. The extra term **S**(*x*) distinguishes Eqn. (7), the LEMUR model, from its special case for **S** ≡ **I**, multicondition PCA, Eqn. (4). To allow translations, we also change *γ*^*′*^(*x*) = *γ*(*x*) + **R**(**X**_*c*:_)*s*_0_(*x*), with *s*_0_ defined below.

Next, we describe the selection of **S** and *s*_0_. It is designed to enable the analyst to state preferences which sets of cells from different conditions should be considered matching each other, i.e., are intended to be the same. We expect such a specification in the form of *E* ∈ ℕ_0_ sets 𝔼_1_, …, 𝔼_*E*_, where each 𝔼_*i*_ ⊂ {1, …, *C*} and 𝔼_*i*_ ∩ 𝔼_*j*_ = ∅ for *i* ≠ *j*. These can be derived, for example, from a set of matching cell type annotations (landmarks) or Harmony’s maximum diversity clustering. This provision of preferences is optional; if it is lacking, *E* = 0, the first term in expression (8) vanishes, the optimization simply results in **S** = **I**, the identity, and LEMUR reverts to multi-condition PCA. **S** is obtained as a solution to the optimization problem

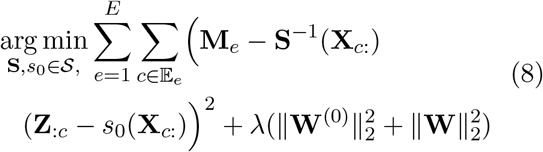

where the optimization domain *𝒮* is described in the next paragraph, and 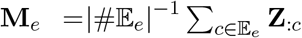 is the mean latent space coordinate of the cells in similarity set 𝔼_*e*_.

The optimization domain *𝒮*, that is, the set of possible **S**(*x*) and *s*_0_, is the set of affine transformations

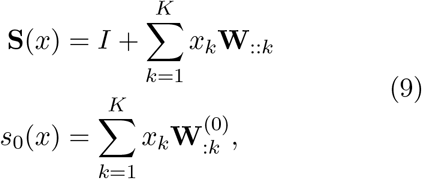

which is parameterized by the 3-tensor **W** with dimensions *P×P ×K* and the *P×K* matrix **W**^(0)^.

The parameter *λ* regularizes the optimization and pulls the result towards **S**(*x*) = **I** and *s*_0_(*x*) = 0.

We provide additional details on how the LEMUR model is implemented in the Supplementary Notes.

### Software details

The analysis was conduced using R version 4.2.2 and Bioconductor version 3.16. We used scVI version 1.1.2, CPA version 0.8.3, Harmony version 1.1.0 and, miloDE version 0.0.0.9000 (git commit id: 8803302).

In the performance assessment, we ran LEMUR with 30 latent dimensions and a test fraction of 0% for Fig. 3A-D and 50% for Fig. 3EG. For the biological vignettes, we used latent space dimensions *P* = 60, 80, and 30 for the glioblastoma, zebrafish, and Alzheimer plaque analyses, respectively.

We provide additional details on software arguments, simulation settings, and benchmark metrics in the Supplementary Notes.

## Supplementary Figures

**Suppl. Figure S1:**
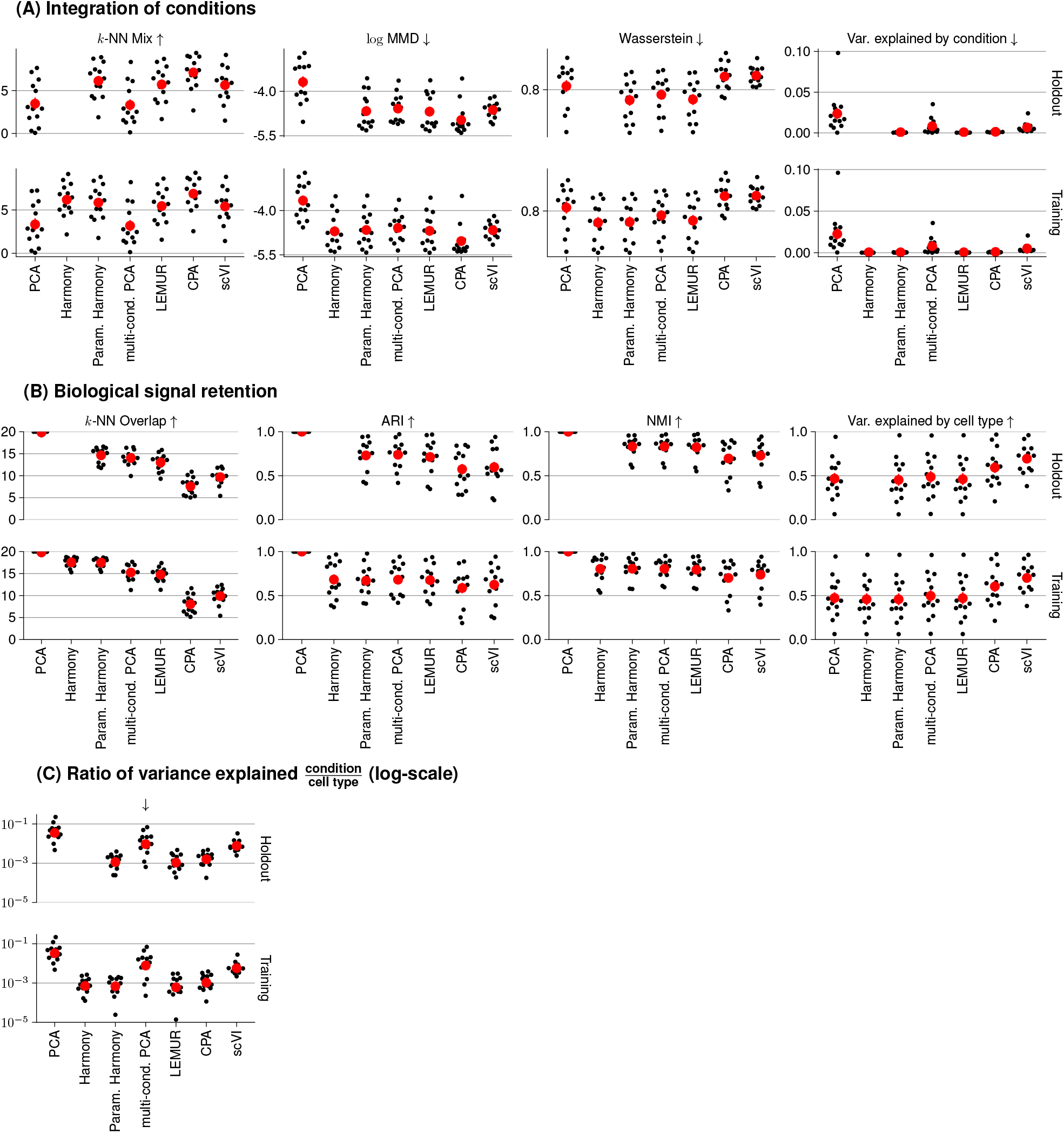
Comparison of integration performance across 13 datasets. (A) Beeswarm plots for each metric comparing how well each method adjusted the latent embedding for the known covariates. The arrows next to the metric indicate if higher or lower values indicate better performance. (B) Beeswarm plots for each metric comparing how well each method retained the biological signal. (C) A beeswarm plot of an integrated performance measure comparing the ratio of variance explained by the known covariates vs cell types (as a proxy for the biological signal). Each black point is the result one dataset and the red points show mean performance. *k*-NN: *k* nearest neighbors, MMD: maximum mean discrepancy, var. expl.: variance explained, ARI: adjusted Rand index, NMI: normalized mutual information.

**Suppl. Figure S2:**
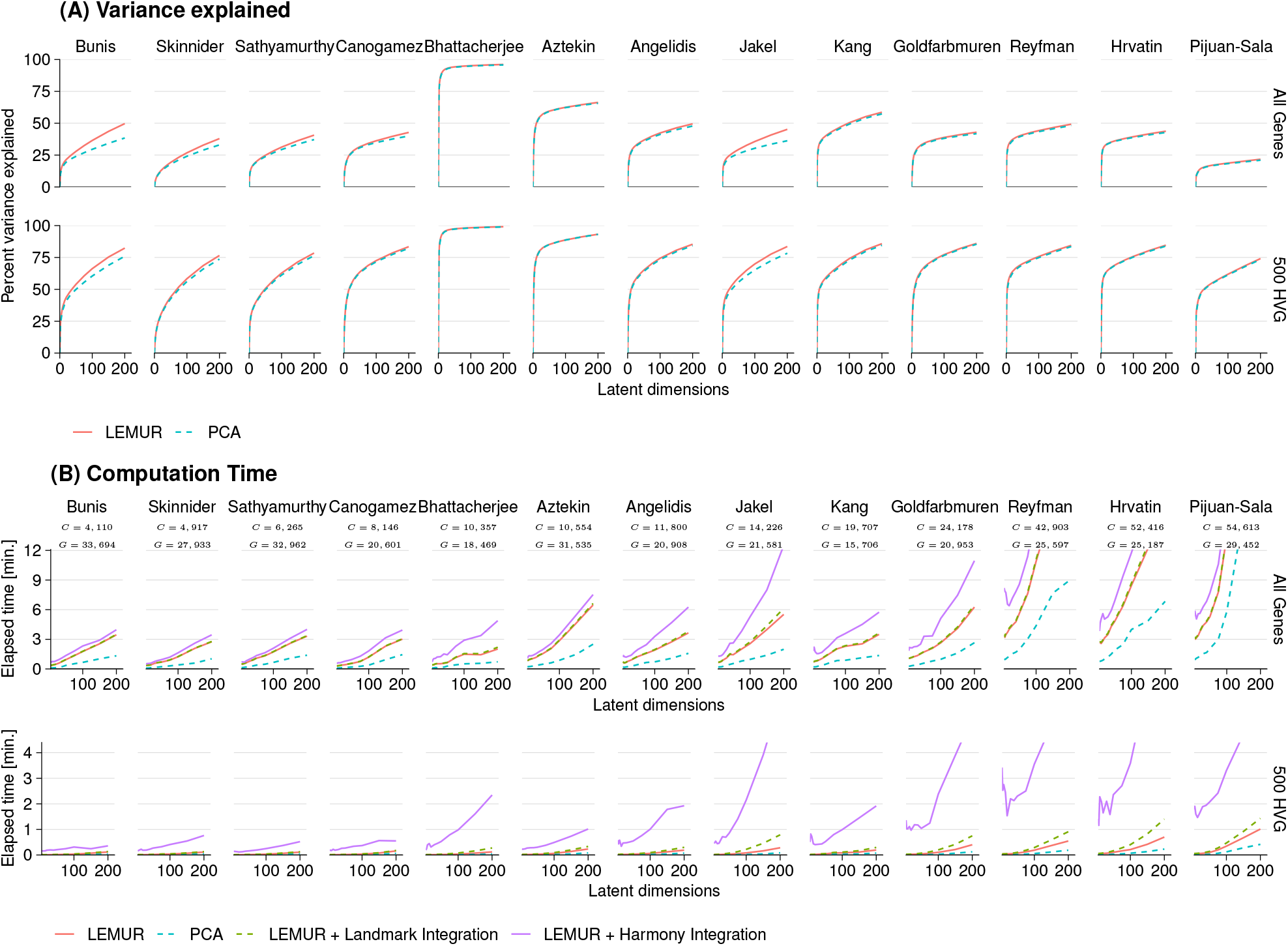
(A) Line plot of number of latent dimensions against the variance explained for PCA and LEMUR. (B) Line plot of number of latent dimensions against the computation time for PCA, LEMUR, and LEMUR with landmark or Harmony-based integration.

**Suppl. Figure S3:**
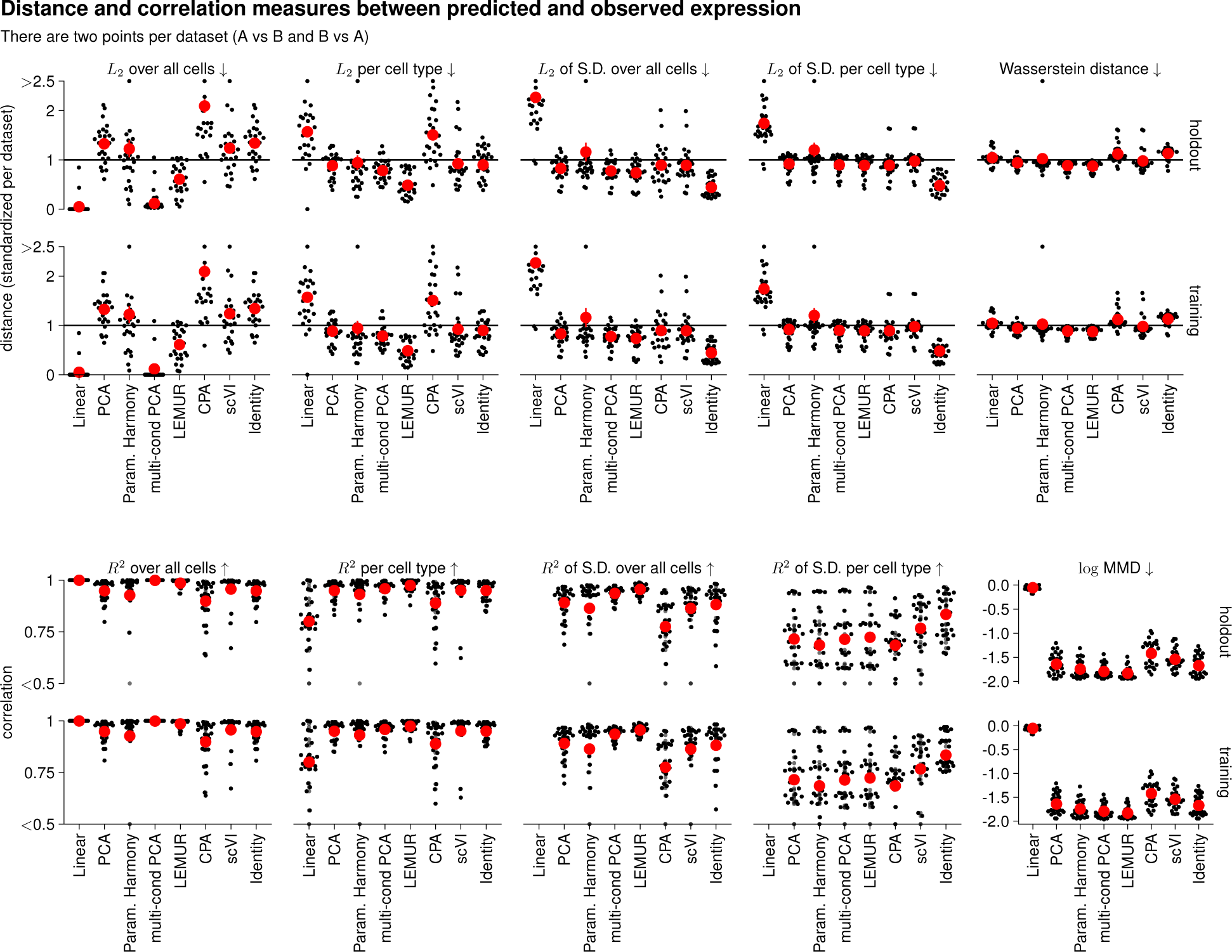
Comparison of the ability of each method to predict the expression of genes across conditions for 13 datasets. The panels show different distance and correlation measures comparing the predicted expression in condition *B* for cells from condition *A* against the observed expression of cells in condition *B* and *vice versa*. The *L*_2_ distance and the correleation where calculated using the mean of the predictions and observations over all cells or per cell type. As the distances varied by two orders of magnitude between datasets, we divided each distance by the mean per dataset. The red points show the mean per method. S.D.: standard deviation, MMD: maximum mean discrepancy.

**Suppl. Figure S4:**
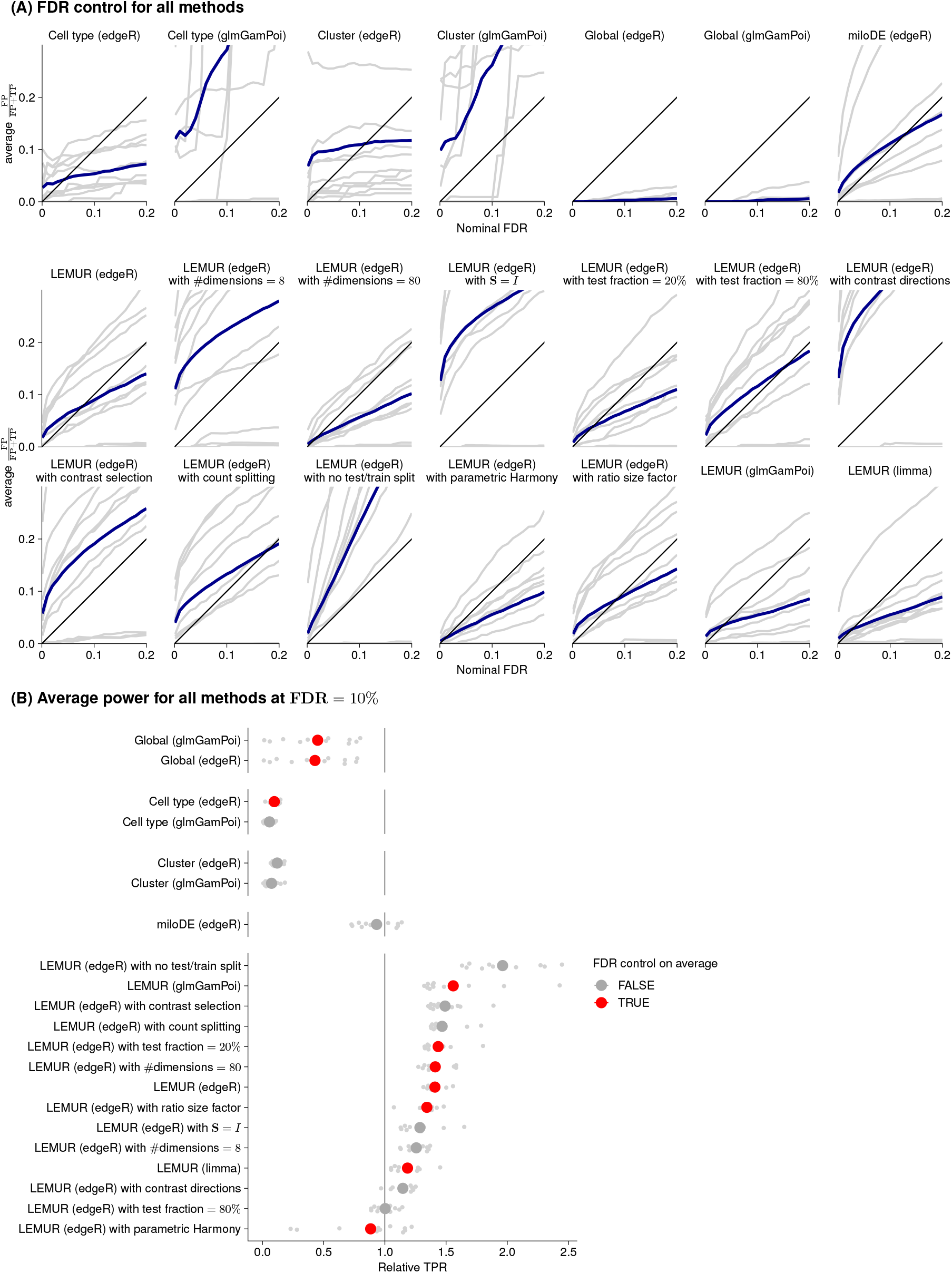
FDR control and power for all methods across eleven datasets where we average across the ten replicates per dataset. (A) Line plot of the nominal FDR against the observed false discovery proportion. The FDR is the expectation of the false discovery proportion over many samples. The blue line shows the average. (B) Scatter plot of the relative TPR at an FDR = 10% for each method across eleven datasets standardized by the mean per dataset. The point range shows the mean and standard error. If the average observed FDR is larger than 10% the point is greyed out. FDR: false discovery rate, TPR: true discovery rate

**Suppl. Figure S5:**
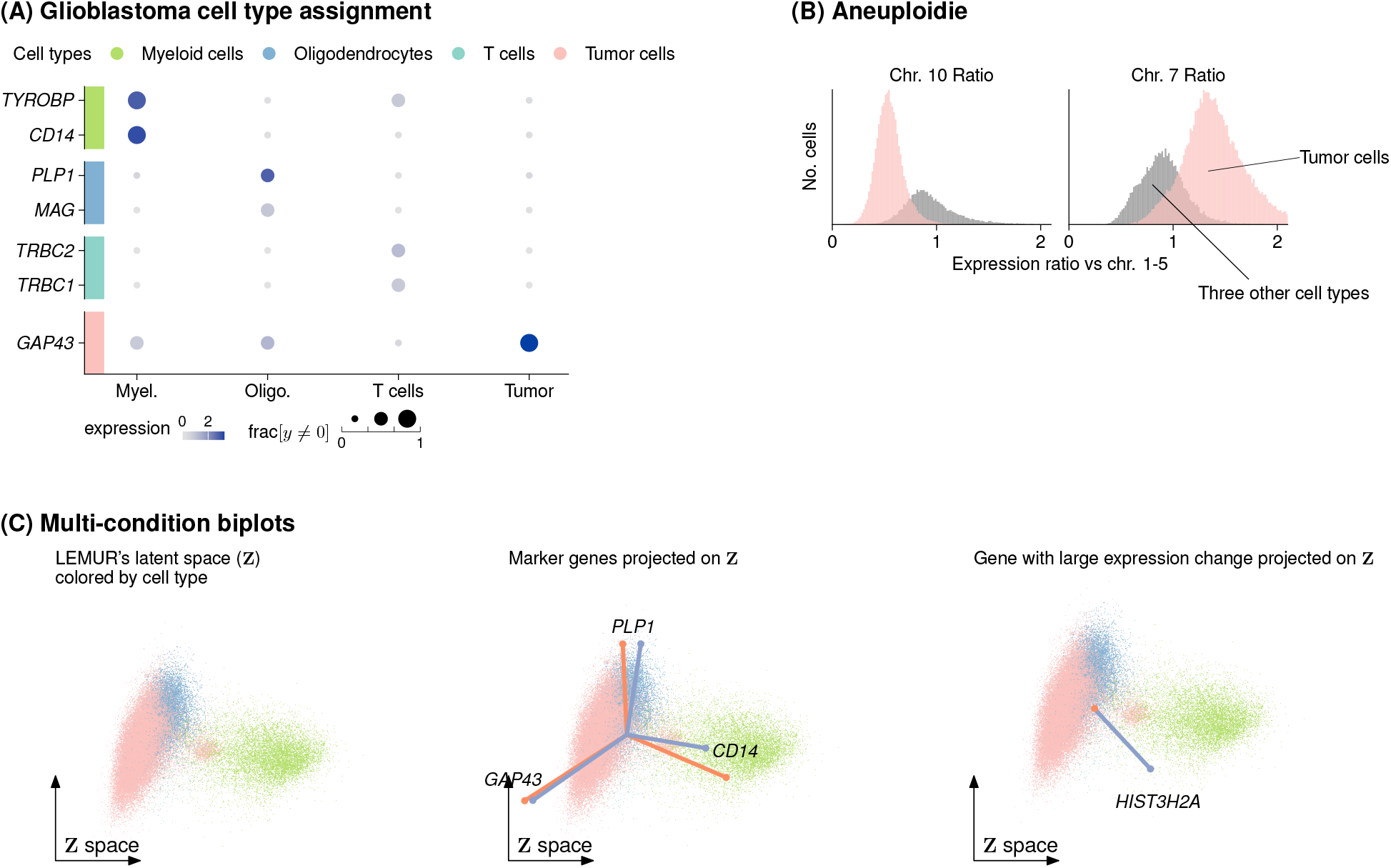
Glioblastoma cell type assignment and multi-condition biplots. (A) Marker gene and (B) chromosome-aggregated expression levels for each cell type. The the tumor cells are known to bear an amplification of chromosome 7 and a deletion of chromosome 10 (Zhao et al., 2021), so we identified the tumor cluster using the ratio of average gene expression on chromosome 7, resp. 10, over the average expression on chromosomes 1-5 (C) Multi-condition biplots showing (left) the first two dimensions of the LEMUR latent space (**Z**) for all cells overlayed with arrows representing (middle) three cell type marker genes from Panel A and (right) a gene with large expression change specifically in myeloid cells (*HIST3H2A*, more details in Suppl. Fig. S6).

**Suppl. Figure S6:**
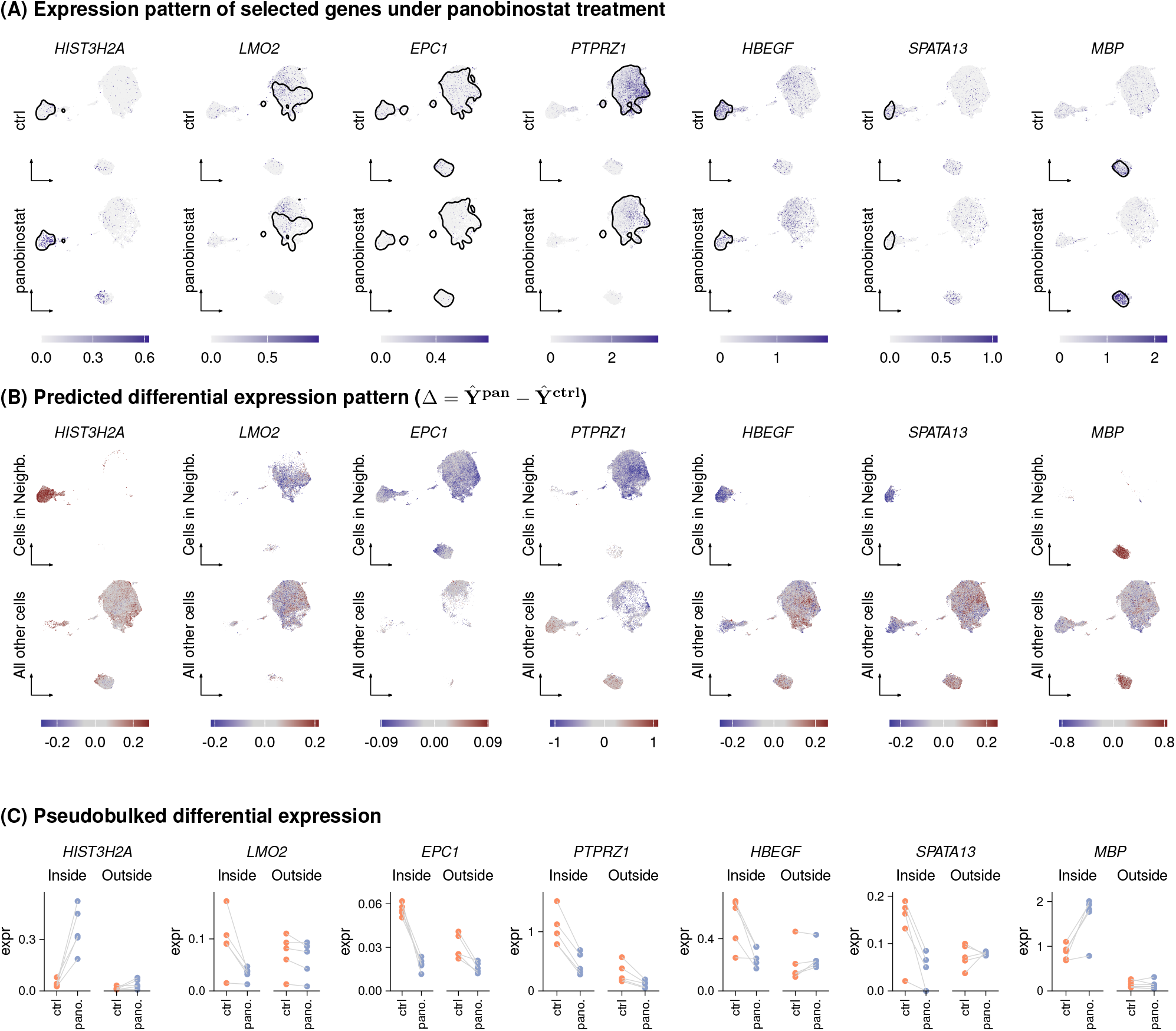
Differential expression patterns for seven genes with the neighborhoods inferred by LEMUR. (A) UMAPs colored by gene expression (log-normalized counts). The black line encircles 80% of the cells inside the gene-specific differential expression neighborhood. (B) UMAPs colored by predicted expression change per cell. The cells are separated depending if they are inside or outside the neighborhood. (C) Scatter plot of the pseudobulked expression values per condition, neighborhood status and sample.

**Suppl. Figure S7:**
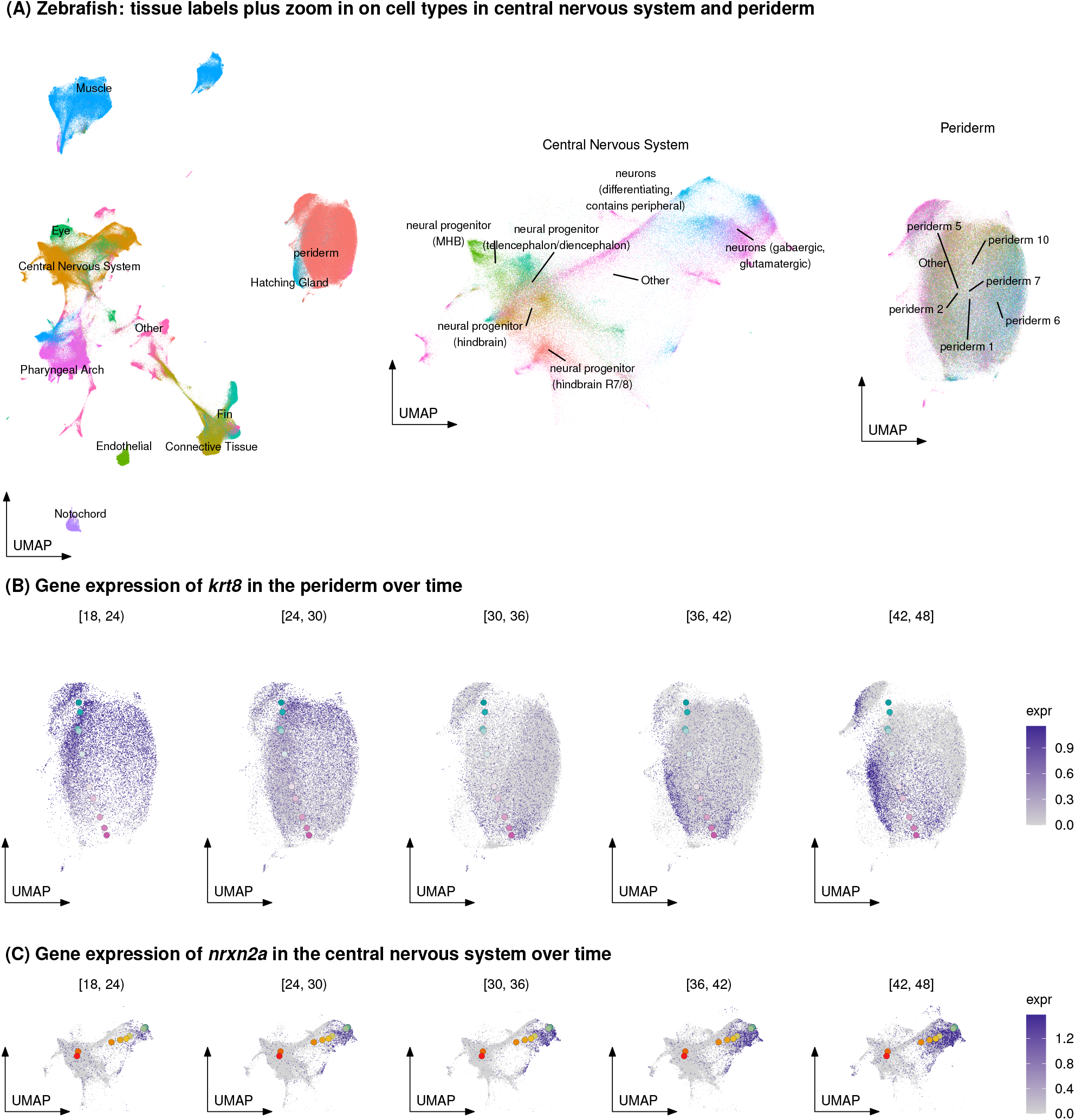
Cell type annotation for zebrafish. (A) Left panel: UMAP of the full timecourse data with cells colored by tissue type as annotated in Saunders et al. (2023). Middle and right panel: Cell type annotations for central nervous system and periderm also from Saunders et al. (2023). (B) Gene expression of *krt8* in the periderm over time. (C) Gene expression of *nrxn2a* in the central nervous system over time. The overlayed points are the synthetic cells from Fig. 5B.

**Suppl. Figure S8:**
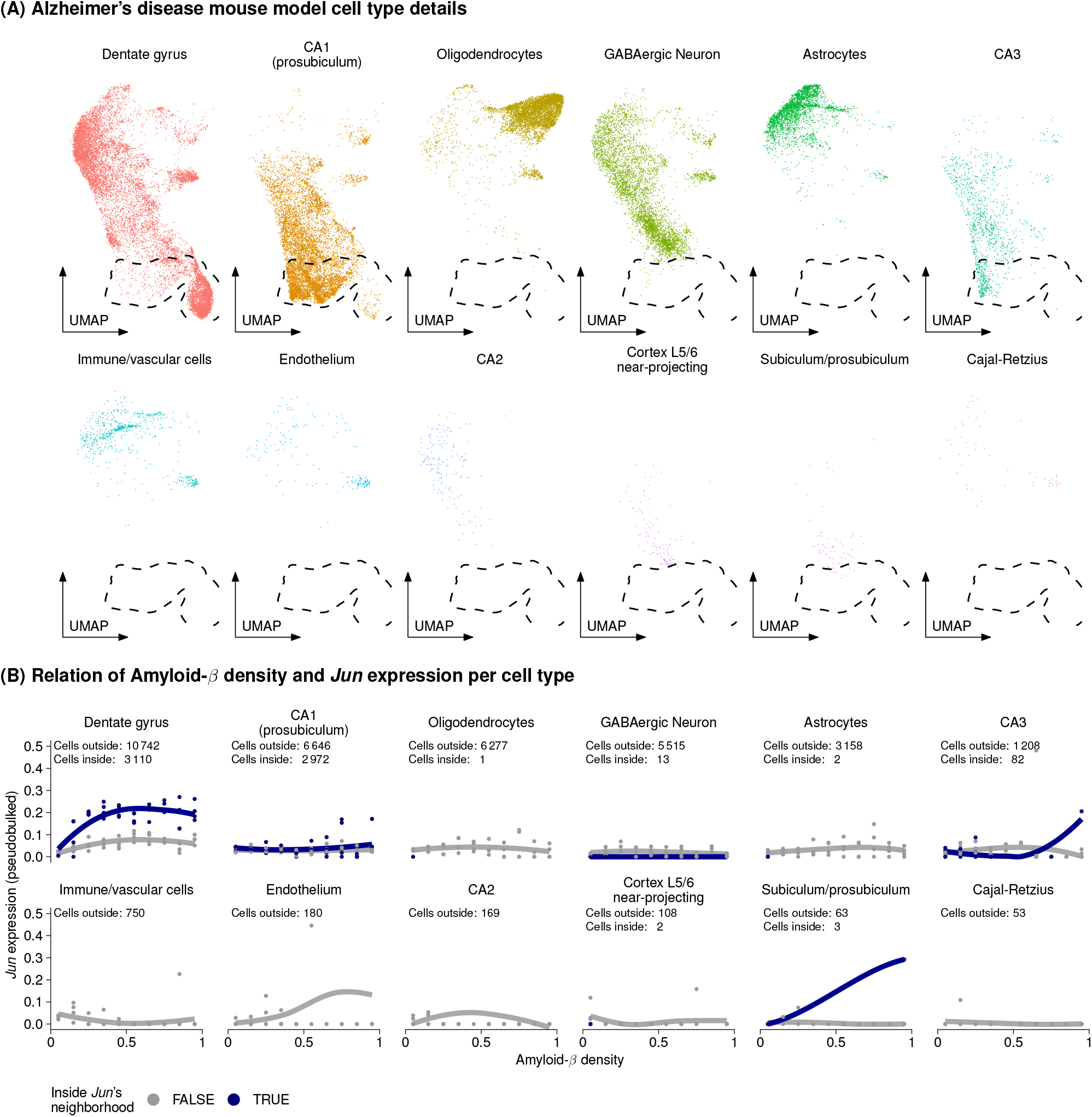
Cell type annotation for mouse brain datasets. (A) UMAP of the Alzheimer’s disease mouse model dataset colored and split by cell type annotation provided by Yao et al. (2021). The dashed shape is the 97% density contour of the cells inside the neighborhood. (B) Scatter plot with smoothing fit of the pseudobulked gene expression of *Jun* against the binned amyloid-*β* plaque density. Each dot is the pseudobulked expression per mouse, cell type, plaque density bin, and neighborhood status. The text at the top of the graph lists the number of cells from that cell type which were inside and outside of the *Jun* neighborhood.

**Suppl. Table S1:** Overview of the patients from whom the glioblastoma biopsies originated.

| Patient ID | Age | Gender | Tumor location | Conditon | # Cells |
| --- | --- | --- | --- | --- | --- |
| PW030 | 65 | M | right parietal | vehicle (DMSO) | 17384 |
|  |  |  |  | 0.2 uM panobinostat | 5686 |
| PW032 | 61 | M | left frontal | vehicle (DMSO) | 5003 |
|  |  |  |  | 0.2 uM panobinostat | 738 |
| PW034 | 68 | F | left parieto-occipital | vehicle (DMSO) | 12375 |
|  |  |  |  | 0.2 uM panobinostat | 1761 |
| PW036 | 56 | M | right temporal | vehicle (DMSO) | 10292 |
|  |  |  |  | 0.2 uM panobinostat | 2558 |
| PW040 | 69 | M | right temporal | vehicle (DMSO) | 8901 |
|  |  |  |  | 0.2 uM panobinostat | 1257 |

## Supplementary Notes

### Implementation

We fit multi-condition PCA, Eqn. (4) by:

1. Solving the linear regression for **Γ**, treating **R**(*x*) and **Z** as 0.
2. Optimizing on the Grassmann manifold for the parameters **B** of the function **R**, keeping Γ fixed.
3. Inferring **Z**_:*c*_ by projecting **Y**_:*c*_ on the orthonormal basis **R**(**X**_:*c*_).

In Step 1, we solve a linear regression for **Γ**

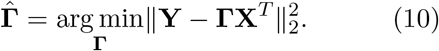

In Step 2, we need to solve the manifold regression problem

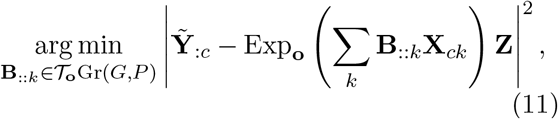

where 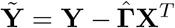. We choose the base point **o** as the first *P* principal vectors of fitting PCA to full, centered data matrix 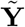.

To optimize **B**, we build on the work of Kim et al. (2014). They developed an algorithm to approximate the geodesic regression problem

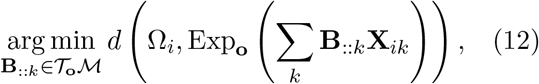

where *ℳ* is a Riemannian manifold, Ω_*i*_ ∈ *ℳ* are data points on the manifolds, and

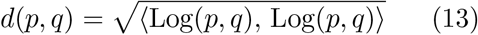

is the geodesic distance between two points on *ℳ*. Here, Log is the inverse of the exponential map.

If the observations Ω_*i*_ are close to each other, the solution to Eqn. (12) is well approximated by the solution to a standard linear regression in the tangent space

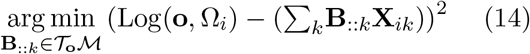

for a base point *o* ∈ *ℳ* that is close to the center of all Ω_*i*_.

Another step is required before we can apply Kim et al. (2014)’s algorithm, since in our case, the observations 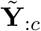 are not elements of the manifold (in our case, *ℳ* is the Grassmann manifold Gr(*G, P*)). We partition {1, …, *C*}, the set of all cells, into sets of cells that share the same condition: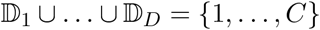 and 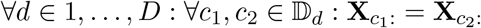. Then, for each *d* (i.e., for each group of cells under the same conditions) we find an orthonormal basis **U**_*d*_ ∈ Gr(*G, P*) using PCA on 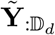, the data for these cells only. We then approximate a solution of Eqn. (11) by linear regression weighted by the number of observations per condition (#?_*d*_) on the *U*_*d*_ projected into the tangent space of **o**. For this, we plug Ω_*i*_ = **U**_*d*_ into Eqn. (14) with

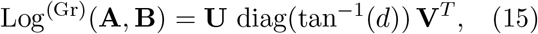

where **U, V**, and *d* come from an SVD of (**B** − **AA**^*T*^ **B**)(**A**^*T*^ **B**)^−1^ = **U** diag(*d*)**V**^*T*^ (Absil et al., 2004).

### Non-distance preserving extension

After fitting the model described in the previous section, the user can choose to also fit a non-distance preserving extensions. The non-distance preserving term **S** is defined by the the parameters of **W** and **W**^(0)^, which we fit using ridge regression with a user-specified penalty that defaults to *λ* = 0.01.

### Post-processing

After fitting the LEMUR model, we adjust the base space so that the rows of **Z** are sorted in descending order of their variance, i.e., we take our specific set of basis vectors and adjust them so that they may be interpreted analogously to principal components, as pointing in the direction of highest, second-highest, … variance. Specifically, we calculate a singular value decomposition of **Z**

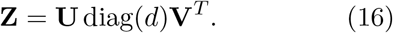

We then set the base point to

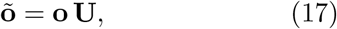

adjust the coefficients of **R** to

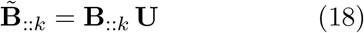

and set the low-dimensional embedding **Z** to

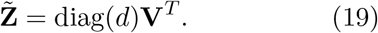

**Table 1.**
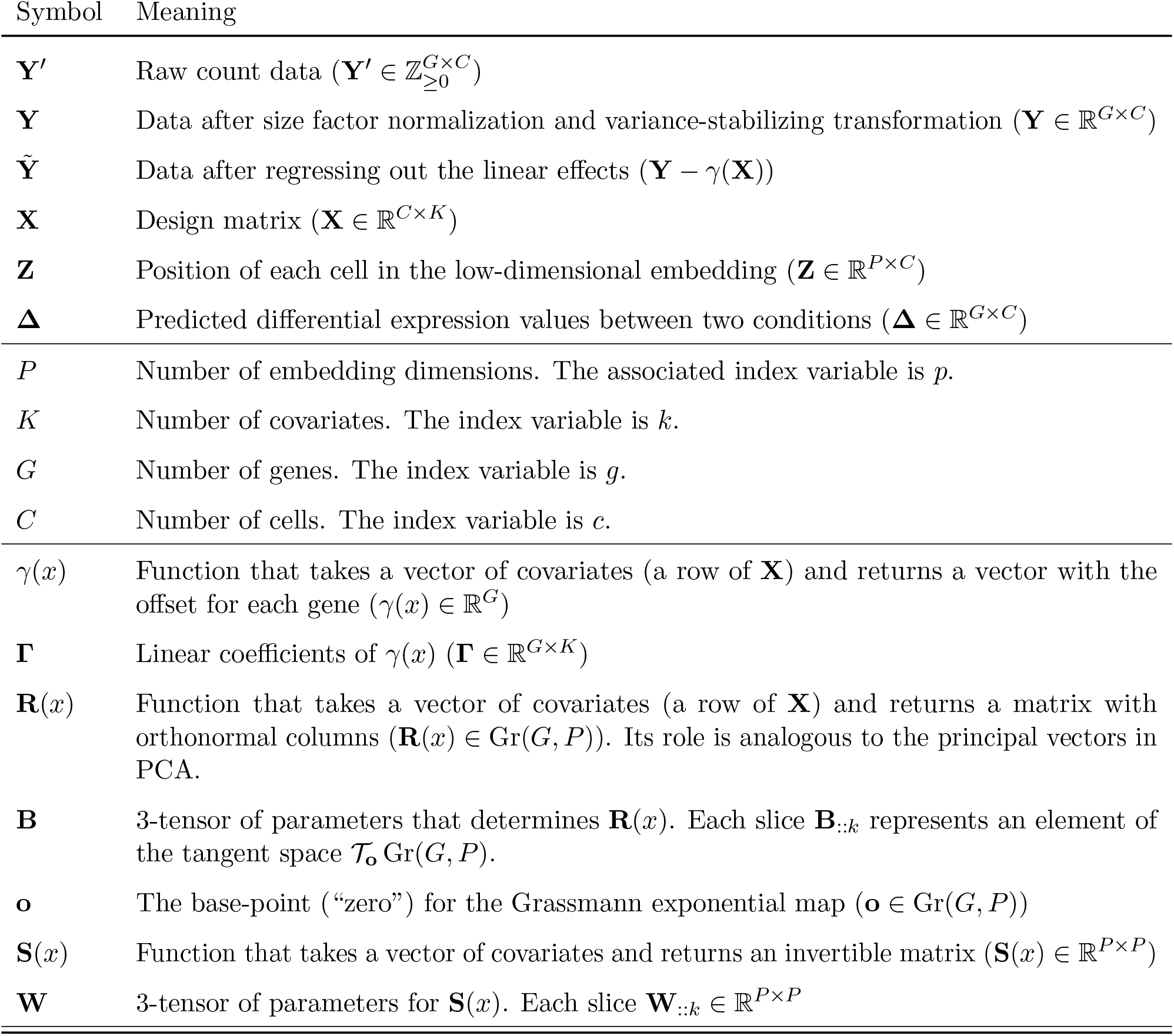
Notation used in the manuscript.

### Cluster-free differential expression

The parametric model of the multi-condition single-cell data learned by LEMUR can be used for intra- and extrapolation, that is, for prediction. First, for any observed cell, it predicts its gene expression in any of the conditions, even though each cell was only observed exactly once. Second, as each cell is parameterized by a position in latent space, the model can also predict expression of “synthetic cells” at any (unobserved) position *z* in the latent space for any condition *x*. We use these capabilities for differential expression analysis. Using the inferred parameters for *γ*(*x*), **R**(*x*), and **S**(*x*), we write

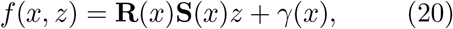

where *f* is a function that predicts the gene expression of a cell at latent space position *z* in the embedding space for any condition *x*.

Thus, the predicted differential expression for all genes in cell *c* between conditions 1 and 2 (*d*_1_, *d*_2_ ∈ ℝ^*K*^) is

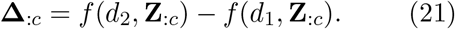

## Differential expression neighborhoods

We use a stochastic sampling algorithm that works on the differential expression matrix **Δ** to identify *neighborhoods* of cells that show consistent differential expression. Intuitively, these are intended to be sets of cells that cluster together in latent space, i.e., are similar or related cell types and cell states, that also show consistent differential expression with respect to the contrast of interest.

In a first step, we sample many onedimensional projections of the embedding **Z**. Specifically, we repeat the following *N* times: randomly sample two cells from {1, …, *C*}, compute the vector between them, 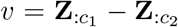, and project the data from all cells onto *v*. This results in the matrix (*w*_*cn*_), where *n* = 1, …, *N* indexes the random samples and *c* indexes the cells. We choose *N* large enough that there is a good chance that interesting differential expression patterns are apparent in one or more of the *w*_:*n*_’s.

Next, for each gene *g*, we identify the best of these one-dimensional data projections by choosing an *n*_*g*_ for which 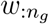 has the maximal absolute correlation to **Δ**_*g*:_. Intuitively, this selects a direction in latent space along which there is differential expression for that gene. We compute the order statistic (*i*_1_, …, *i*_*C*_) of 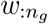, for each *c* = 1, …, *C* compute the *t*-statistic for the sample 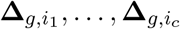, choose the *c*_*g*_ which maximizes that statistic, and set 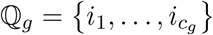.

## Pseudobulk differential expression analysis

The pseudobulk approach accounts for the fact that the most relevant unit of replication in multicondition single-cell data is the sample (and not the cells) (Crowell et al., 2020). It works by summing the counts across cells within a sample from the same cell subpopulation (e.g., “cell type”). If the data were obtained from *F* samples, the information which cell belongs to which sample implies a partition of {1, …, *C*} into *F* sets of cells, which we call 𝔽_1_, …, 𝔽_*F*_.

Let 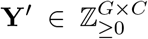 be the count matrix from which **Y** was constructed. Then we form the pseudobulk count matrix **V** ∈ ℤ^*G×F*^ as

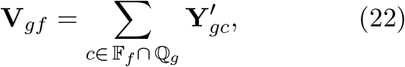

for *f* = 1, …, *F*, and calculate a gene-specific size factor

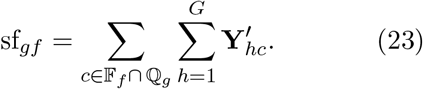

This construction differs from the usual pseudobulk approach as it uses a different set of cells ℚ_*g*_ for each gene.

To address the post-selection inference problem (we want to make inference on the statistical significance of an observed trend based on a test statistic that was itself constructed after consulting the data), we compute (22) on a set of held out cells, which are matched via the (*w*_*cn*_) matrix. The fraction of cells that we assign to the training set (which is used to compute the neighborhoods ℚ_1_, …, ℚ_*G*_), and to the test set (which is used to compute the pseudobulk data (22)) balances the power for accurately identifying neighborhoods with interesting gene expression changes, versus the power to provide statistical significance statements for those identifications. Future work may find more elegant solutions.

## Execution details

### Integration and prediction benchmark

The integration benchmark measured the ability of methods to adjust for known covariates while retaining the biological structure of the data. The prediction benchmark measured for the tools which support it how well they are able to predict the expression of a cell in arbitrary conditions.

For both benchmarks, we used 13 single-cell datasets (Angelidis et al., 2019; Aztekin et al., 2019; Bhattacherjee et al., 2019; Bunis et al., 2021; Cano-Gamez et al., 2020; Goldfarbmuren et al., 2020; Hrvatin et al., 2018; Jäkel et al., 2019; Kang et al., 2018; Pijuan-Sala et al., 2019; Reyfman et al., 2019; Sathyamurthy et al., 2018; Skinnider et al., 2021) which we downloaded from publicly available sources (see Data Availability for details).

We preprocessed each dataset using transformGamPoi package’s shifted log transform function with the default parameters. We identified the 500 most variable genes and held out 20% randomly chosen cells.

For the integration benchmark, we compared scVI version 1.1.2 (Lopez et al., 2018), CPA version 0.8.3 (Lotfollahi et al., 2023), Harmony version 1.1.0 (Korsunsky et al., 2019), LEMUR with *S*≡ *I* (multi-condition PCA), LEMUR with R fixed to the principal vectors of 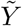 (parametric Harmony), and the full LEMUR model. For PCA, we used the fast implementation from the irlba package (version 2.3.5.1).

For the prediction benchmark, we did not consider Harmony, as it does not support going back from the integrated embedding to the gene expression space. Instead, we included two other comparisons, which can be considered baseline controls: a linear model-based method that predicts the mean of each condition, and *identity prediction*, which returns the original expression observed for a cell independent of the requested condition.

We tried to run the methods as much as possible with the default parameters. On the advice of the authors, we ran CPA directly on the counts (that is, not on the variance stabilized data) with the parameters from the tutorial on integrating the Kang dataset (most importantly, 64 latent dimensions, a negative binomial loss, and a learning rate of 0.0003). For the predictions from CPA, we, again based on the recommendations of the authors, log-transformed the predicted counts before comparing them to the variance stabilized data. For LEMUR, we used 30 latent dimensions and a test fraction of 0%. We ran PCA with 30 latent dimensions.

For the evaluation of the integration results displayed in Fig. 3B, we used the integration performance of the held out data compared to the training data, except for Harmony, which only integrates its training data and has no concept of integrating further, previously unseen data. To make the output of the different methods comparable, we brought each embedding to a common scale by subtracting the mean for each latent dimension and dividing by the average cell vector length. On this rescaled input, we calculated nine different metrics. Four metrics were used to assess the adjustment for the known covariates:

#### k nearest neighbor (k-NN) mixing

We identified the *k* = 20 nearest neighbors from the training data for each cell from the held out data. We then calculated how many of those 20 neighbors were from the same condition as the original cell. We averaged these values across all held out cells to derive a single metric for each dataset and method pair.

#### Maximum mean discrepancy (MMD)

We calculated the MMD discrepancy with a radial basis function kernel between the held out and training data after subsampling both to a common number of cells (Gretton et al., 2012). We calculated the discrepancy using scaling factors between 10 and 10^−3^ (50 values, log spaced) and 100, 200, and 500 cells and finally averaged the results to get a single metric.

#### Wasserstein distance

We calculated the Wasserstein distance between held out and training cells using the wasserstein function from the transport package after subsampling to a common number of cells. We averaged the results for 100, 200, and 500 cells.

#### Variance explained by condition

We calculated the ratio of residual variance after accounting for known covariates over the total variance of the embedding.

We used these four metrics to assess how well each embedding retained the biological information:

#### kNN overlap

We calculated a reference embedding with PCA, scVI, and CPA that did not try to integrate the conditions. Then we compared for each condition the similarity of the nearest neighbors on the integrated and non-integrated embeddings. We used the PCA as a reference for Harmony and LEMUR. CPA did not support fitting an embedding without a condition variable, thus we randomly assigned each cell to a condition and thus created a perfectly mixed dataset where no additional integration was needed.

#### Adjusted Rand index (ARI)

We calculated a reference embedding with PCA, scVI, and CPA that did not try to integrate the conditions. Then we compared the similarity of a walktrap clustering on the integrated and non-integrated embeddings. We calculated the adjusted Rand index to measure the cluster consistency using the ARI function from the aricode package.

#### Normalized mutual information (NMI)

We followed the same procedure as for the ARI, but calculated the normalized mutual information using the NMI function.

#### Variance explained by cell type

We calculated the ratio of residual variance after accounting for the cell types as annotated in the original data over the total variance of the embedding.

Lastly, we also considered one merged metric that directly contrasts the adjustment for the known covariates and the retention of the biological information

#### Variance explained by condition vs. cell type

We calculated the ratio of residual variance after accounting for the known conditions plus the cell types over the residual variance accounting only for the cell types.

For the prediction benchmark, we considered a total of ten metrics. We considered two conditions for each dataset and always calculated the predicted expression for condition B for cells from condition A against the observed expression in condition B and vice versa.

#### L_2_ mean

The *L*_2_ distance between the mean prediction against the mean observed expression across the whole data.

#### L_2_ mean per cell type

The *L*_2_ distance between the mean prediction against the mean observed expression for each cell type.

#### L_2_ of the standard deviation (S.D.)

The *L*_2_ distance between the standard deviation of the prediction against the standard deviation of the observed expression across the whole data.

#### L_2_ of the S.D. per cell type

The *L*_2_ distance between the standard deviation of the prediction against the standard deviation of the observed expression for each cell type.

#### R^2^ mean

The correlation between the mean prediction against the mean observed expression across the whole data.

#### R^2^ mean per cell type

The correlation between the mean prediction against the mean observed expression for each cell type.

#### R^2^ of the S.D

The correlation between the standard deviation of the prediction against the standard deviation of the observed expression across the whole data.

*R*^2^ *of the S.D. per cell type*

The correlation between the standard deviation of the prediction against the standard deviation of the observed expression for each cell type.

#### Maximum mean discrepancy (MMD)

We calculated the MMD discrepancy between the predicted and observed data after subsampling to a common number of cells with a radial basis function kernel (Gretton et al., 2012). We calculated the discrepancy using scaling factors between 10 and 10^−3^ (50 values, log spaced) and 100, 200, and 500 cells and finally averaged the results to get a single metric.

#### Wasserstein distance

We calculated the Wasserstein distance between predicted and observed data using the wasserstein function from the transport package after subsampling to a common number of cells. We averaged the results for 100, 200, and 500 cells.

## Variance explained comparison

For all 13 datasets used in the integration and prediction benchmark, we compared the fraction of variance explained by LEMUR and by PCA. To make the results comparable, we manually regressed out the effects of the known covariates. We compared irlba’s approximate PCA implementation and the LEMUR model accounting for the known covariates. We fixed the linear coefficient estimator =“zero” as we had already manually removed the linear effects. We measured the elapsed time using R’s system.time function and the memory using the GNU time command.

## Differential expression and neighborhood inference benchmark

For the differential expression benchmark, we took the gene expression values of the top 8000 highly variable genes from individual conditions and appended 200 simulated genes. We assigned each cell to a synthetic control or treatment condition. This ensured that for all original genes there was no real differential expression, whereas for the 200 simulated genes we were able to control the number of cells with a differential expression pattern and the log fold change.

We only considered the dataset-condition combinations which had the most independent replicates. We chose two conditions from the Angelidis et al. (2019); Goldfarbmuren et al. (2020); Kang et al. (2018) data, three conditions from Hrvatin et al. (2018) and one from Sathyamurthy et al. (2018). Thus, we had eleven datasets in total.

We then simulated 200 genes for each dataset with varying number of affected cells, which we repeated ten times per dataset. We first performed *k* means clustering on the 50-dimensional embedding of the data with either 2, 3, 10, or 20 clusters. Then, for each gene, we chose one of the clusters and fixed the log fold change to 0.5, 1, 2, or 4, respectively (i.e., for the smaller clusters, we used a larger effect size). The counts were simulated according to

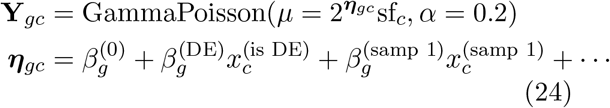

where 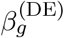 is the log fold change, 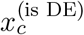 indicates if cell *c* is inside the selected cluster, sf_*c*_ is the size factor for cell *c* calculated on the observed genes, and 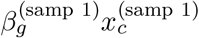 simulates a sample-specific effect where 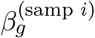 is drawn from a normal distribution with a standard deviation of 0.1.

Using the known set of simulated genes (true positive) and original genes (true negatives), we calculate the true positive rate (TPR, fraction of identified true positives) and false discovery proportion (fraction of false positives among all positives).

Our default settings for LEMUR in the differential expression benchmark were

- 30 latent dimensions,
- a test fraction of 50%,
- directions =“randomized”,
- selection procedure =“zscore”,
- size factor method =“normed sum”,
- and test method=“edgeR”.

In addition, we tested several variations:

- LEMUR with test method set to *glmGamPoi* (Ahlmann-Eltze and Huber, 2020) or *limma* (Smyth, 2004),
- LEMUR with 8 or 80 latent dimensions,
- LEMUR with a test fraction of 20% and 80%,
- LEMUR with size factor method =“ratio”,
- LEMUR with directions =“contrast” or

selection procedure =“contrast”,

- LEMUR with **S** ≡ **I** (multi-condition PCA) or **R** fixed to the principal vectors of 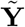(parametric Harmony),
- LEMUR where we reused the training data for testing,
- LEMUR where the test and training data were generated through count splitting (Neufeld et al., 2022).

We compared the FDR and TPR from LEMUR against seven alternative methods:

*Global test* A single test with *edgeR* or *glmGamPoi* across the full dataset.

*Cell type test* One test per cell type (using the annotation from the original data) with *edgeR* or *glmGamPoi*.

*Cluster test* One test per walktrap cluster on the Harmony integrated data with *edgeR* or *glmGamPoi*.

*MiloDE* One test per Milo neighborhood using *edgeR* (Missarova et al., 2024).

For the *cell type test, cluster test*, and MiloDE, which perform more than one test per gene, we considered a group of cells as positive if they contained more than 60% changed cells (at least 10). If none of the cell groups for a gene fulfilled this criterion, the group with the largest fraction of changed cells was considered as positive. A group of cells was considered negative if less than 10% of the cells were changed. If the fraction of changed cells was between 10% and 60% the status for that group of cells was considered indeterminate and the group was ignored for the TPR and FDR calculation.

## Glioblastoma analysis

We downloaded the count data and patient annotations from GSE148842. We transformed the counts using the function shifted log transform from the transformGamPoi package and filtered out all genes with a total of less than 6 counts. We filtered out all cells which did not pass the quality filters from the scuttle package’s perCellQCFilters function, and also those cells that had less than 800 or more than 12 000 counts.

We assigned the cell types and tumor status following the methods from the original publication (Zhao et al., 2021). We clustered the Harmony integrated data with walktrap clustering into four clusters. Zhao et al. (2021) identified a chromosome 7 duplication and 10 deletion in all samples. Accordingly, we assigned the cluster which showed upregulation of genes from chromosome 7 and downregulation of genes from chromosome 10 as the tumor cells. The other three clusters were assigned based on marker genes as shown in Suppl. Fig. S5A.

We ran LEMUR with 60 latent dimensions and a test fraction of 50%; otherwise, we used the same defaults as in the differential expression benchmark. The full dataset consisted of three conditions: control, panobinostat, and etoposide.

For the analysis, we decided to focus on the contrast between panobinostat and control and do not show any data from etoposide-treated cells.

The color scale of **Δ** in Fig. 4C was capped at the 95% quantile of the absolute values, squishing more extreme values to the range. The difference of difference test in the rightmost panel of Fig. 4C was significant at FDR *<* 0.1 considering all genes with a significant difference (FDR *<* 0.1) between control and panobinostat inside the neighborhood.

We identified gene ontology terms related with up and down-regulated genes using the clusterProfiler package’s enrichGO function on the 200 genes with the smallest p-value comparing them against the universe of genes that passed quality control.

## Zebrafish embryonic development analysis

The data downloaded from GSE202639 was already quality-controlled. We subsetted the full dataset to the control cells (*ctrl-inj* and *ctrl-uninj*) for the 16 time points between 18 hours and 48 hours and the 2 000 most variable genes. We transformed the counts using transformGamPoi package’s shifted log transform function and fit a natural spline model with three degrees of freedom and 80 latent dimensions. We tested the difference between the 48 hour and the 18 hour time point using the settings from the differential expression benchmark.

For the interpolation in Fig. 5B, we selected two pairs of cells and linearly interpolated their latent position. The selected cells were not from the same timepoint, but as they only served as anchors in the latent space **Z** this did not influence the results. We projected ten synthetic cells onto the 2-dimensional UMAP. We calculated the mean of the observed expression values from the 50 nearest neighbors to five synthetic cells at interpolation points 0, 0.25, 0.5, 0.75 and 1. We predicted the gene expression according to a spline fit for all ten synthetic cells at 50 time points equally spaced between 18 and 48 hours.

To prioritize the genes that we manually inspected, we tested whether a spline model with 5 degrees of freedom could significantly better explain the observed expression pattern over time than a linear model within a selected cell type.

## Alzheimer plaque spatial analysis

We downloaded the expression data for the four mouse hippocampi with the Alzheimer plaque densities from the BROAD’s single-cell repository. We subsetted the genes to a common set and filtered out lowly expressed genes (total counts per gene less than 50). We further filtered out cells with more than 20% mitochondrial reads and less than 200 or more than 5 000 total counts.

We fit LEMUR with 30 latent dimensions and a test fraction of 60% on an ordered factor of the plaque density cut into ten equally sized bins. We contrasted the largest bin against the smallest bin using the same settings as in the differential expression benchmark.

## Relation to other methods

### Relation to PCA regression and partial least squares regression

For linear regression problems

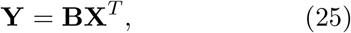

where the design matrix contains many (and potentially some colinear) columns, PCA regression replaces the original design matrix with a lower dimensional approximation **V** produced with PCA:

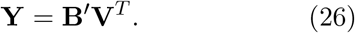

Partial least squares is used in similar circumstances but chooses the lower dimensional approximation **V** such that it is optimal for predicting **Y** (Wold et al., 2001).

The difference to the LEMUR model is that partial least squares and principal component regression find approximations of the design matrix, whereas LEMUR finds low-dimensional subspaces that approximate **Y** and expresses the relation of the subspaces as regression problem.

### Relation to GSFA

Zhou et al. (2023) described a model for the analysis of perturbation single-cell data called *guided sparse factor analysis* (GSFA) which is related to the SupSVD model (Li et al., 2016).

It is built around matrix factorization of the observed data

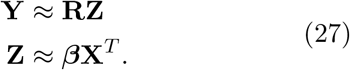

The model uses sparse priors on **R** and ***β*** to identify which gene modules are related to which latent factors. Unlike LEMUR, model (27) makes the latent embedding Z dependent on the design matrix **X**.

The sparsity priors favor situations where a perturbation only affects a small number of latent factors in **Z** and thus columns of **R**. In that case, the contrast between the subspaces spanned by the active factors of **R** and the inactive factors for a perturbation has a similar interpretation as the corresponding comparison in LEMUR. However, LEMUR directly identifies the subspaces per condition and does not need to rely on the indirect effect of the sparsity priors.

### Relation to GEDI

A recent preprint by Madrigal et al. (2023), published after the first preprint of this work (doi: 10.1101/2023.03.06.531268), presents a model for cluster-free differential expression analysis, similar to what we delineate by Step 1 in Fig. 1A.

Their model can be summarized as

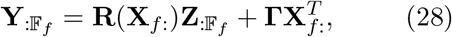

where 𝔽_*f*_ are the indices of all cells from sample *f* and **X** is a design matrix on the sample level (*F × K*). They define the function **R** as the linear combination of a reference state (*θ*_*r*_) and deviations from that reference state (*δ*)

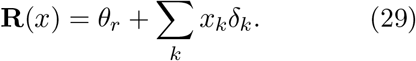

This model is fairly similar to LEMUR as it also works by treating the subspace as a function of the covariates. The biggest difference with our model is that their **R** does not enforce the orthonormality constraints on **R**(*x*). The resulting degeneracy between **R** and **Z** is partially resolved by scaling each row of **Z** to a length of 1 and regularization on **R**, which can optionally be guided by gene-regulatory networks.

### Relation to interaction models

Model (4) can express interactions between known covariates and the latent position of each cell. For example, a drug perturbation might affect the gene expression of cells early in a developmental trajectory more than in mature cells. Our model simultaneously identifies the latent position and the interacting drug effect. Yet, the way the interactions are modeled here differs from that in ordinary linear models.

Interactions in ordinary linear models are formed using a direct (Hadamard) product between two or more known covariates. For example, the effectiveness of trastuzumab on breast cancer cells depends on their HER2 status, i.e., the drug is more effective if the HER2 protein level is high. Accordingly, we could model cell viability as a function of

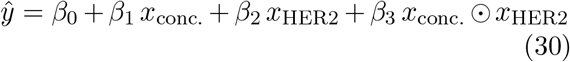

and call *β*_3_ the interaction coefficient.

The LEMUR multi-condition PCA model (4) and the interaction model (30) are closely related.

To demonstrate, we define **R** not as eq. (5) but as

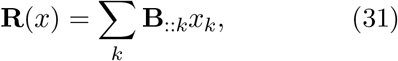

and assume that **X** contains the known drug concentration from our example above, while **Z** contains the latent *HER2* status:

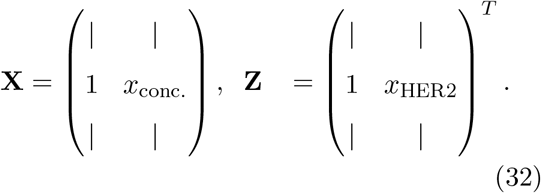

When we plug Eqn. (31) into Ŷ_:*c*_ = **R**(**X**_*c*:_)**Z**_*c*:_, we can rewrite it as

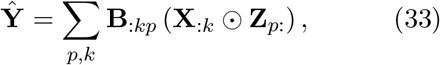

which is just a different way to write eq. (30). This demonstrates that the only difference between the interaction model with latent factors and LEMUR is the choice of **R**.

Interaction model (33) has been used to model the effects of regulatory variants in single-cells across cell states (Cuomo et al., 2022; Nathan et al., 2022). There, the cell states were represented using continuous factors **Z**; however, the estimation proceeded step-wise: first, estimating **Z** using PCA or Harmony and only then fitting the interaction coefficients **B**.

Independent of the parametrization (Eqn. (5) or Eqn. (31)), **R**(*x*) can be interpreted as spanning the space that best approximates the observations from condition *x*. The advantage of Eqn. (5) is that the constraints of the Grassmann manifold naturally map to this intuition. In contrast, the parametrization of Eqn. (31) does not enforce orthonormality between the columns of **R**(*x*), it does not even enforce a common scale. This makes the model degenerate when inferring **Z** and **B** simultaneously.

Geometrically, the columns of **B** in Eqn. (31) that correspond to the intercept in **X** span a base space. All other columns in **B** are vectors that point out of that base space. In contrast, the **B**_::*k*_ ∈ *𝒯* _**o**_Gr(*G, P*) in Eqn. (5) correspond to rotations of the base space. For small angles between the spaces of two conditions, there is little difference between a rotation and the straight vector. Thus, one can interpret our multi-condition PCA model as approximating a conventional interaction model between observed and latent covariates.

## Footnotes

1 We use a recycling convention like the one used in the R language for the sum operator (+) for a matrix *A* and a conformable column vector *b*: (*A* + *b*)_*ij*_ = *A*_*ij*_ + *b*_*i*_

## References

Absil, P.-A., Mahony, R. and Sepulchre, R. (2004). Riemannian geometry of Grassmann manifolds with a view on algorithmic computation. Acta Applicandae Mathematica 80: 199–220, doi: 10.1023/B:ACAP.0000013855.14971.91.

Ahlmann-Eltze, C. and Huber, W. (2020). glmGamPoi: fitting Gamma-Poisson generalized linear models on single cell count data. Bioinformatics 36: 5701–5702, doi: 10.1093/bioinformatics/btaa1009.

Ahlmann-Eltze, C. and Huber, W. (2023). Comparison of transformations for single-cell RNA-seq data. Nature Methods 20: 665–672, doi: 10.1038/s41592-023-01814-1.

Akhter, R., Sanphui, P., Das, H., Saha, P. and Biswas, S. C. (2015). The regulation of p53 up-regulated modulator of apoptosis by JNK/c-Jun pathway in β-amyloid-induced neuron death. Journal of Neurochemistry 134: 1091–1103, doi: 10.1111/jnc.13128.

Angelidis, I., Simon, L. M., Fernandez, I. E., Strunz, M., Mayr, C. H., Greiffo, F. R., Tsitsiridis, G., Ansari, M., Graf, E., Strom, T.-M. et al. (2019). An atlas of the aging lung mapped by single cell transcriptomics and deep tissue proteomics. Nature Communications 10: 963, doi: 10.1038/s41467-019-08831-9.

Aztekin, C., Hiscock, T., Marioni, J., Gurdon, J., Simons, B. and Jullien, J. (2019). Identification of a regeneration-organizing cell in the xenopus tail. Science 364: 653–658, doi: 10.1126/science.aav9996.

Baglama, J., Reichel, L. and Lewis, B. W. (2022). irlba: Fast Truncated Singular Value Decomposition and Principal Components Analysis for Large Dense and Sparse Matrices. doi: 10.32614/CRAN.package.irlba, r package version 2.3.5.1.

Bendokat, T., Zimmermann, R. and Absil, P.-A. (2023). A Grassmann manifold handbook: Basic geometry and computational aspects. arXiv doi: 10.48550/arXiv.2011.13699.

Bhattacherjee, A., Djekidel, M. N., Chen, R., Chen, W., Tuesta, L. M. and Zhang, Y. (2019). Cell type-specific transcriptional programs in mouse prefrontal cortex during adolescence and addiction. Nature Communications 10: 4169, doi: 10.1038/s41467-019-12054-3.

Bunis, D. G., Bronevetsky, Y., Krow-Lucal, E., Bhakta, N. R., Kim, C. C., Nerella, S., Jones, N., Mendoza, V. F., Bryson, Y. J., Gern, J. E. et al. (2021). Single-cell mapping of progressive fetal-to-adult transition in human naive T cells. Cell Reports 34, doi: 10.1016/j.celrep.2020.108573.

Bunne, C., Stark, S. G., Gut, G., Del Castillo, J. S., Levesque, M., Lehmann, K.-V., Pelkmans, L., Krause, A. and Rätsch, G. (2023). Learning single-cell perturbation responses using neural optimal transport. Nature Methods doi: 10.1038/s41592-023-01969-x.

Butler, A., Hoffman, P., Smibert, P., Papalexi, E. and Satija, R. (2018). Integrating single-cell transcriptomic data across different conditions, technologies, and species. Nature Biotechnology 36: 411–420, doi: 10.1038/nbt.4096.

Cable, D. M., Murray, E., Shanmugam, V., Zhang, S., Zou, L. S., Diao, M., Chen, H., Macosko, E. Z., Irizarry, R. A. and Chen, F. (2022). Cell type-specific inference of differential expression in spatial transcriptomics. Nature Methods 19: 1076–1087, doi: 10.1038/s41592-022-01575-3.

Cano-Gamez, E., Soskic, B., Roumeliotis, T. I., So, E., Smyth, D. J., Baldrighi, M., Willé, D., Nakic, N., Esparza-Gordillo, J., Larminie, C. G. et al. (2020). Single-cell transcriptomics identifies an effectorness gradient shaping the response of CD4+ T cells to cytokines. Nature Communications 11: 1801, doi: 10.1038/s41467-020-15543-y.

Chambers, J. and Rabbitts, T. H. (2015). LMO2 at 25 years: a paradigm of chromosomal translocation proteins. Open Biology 5: 150062, doi: 10.1098/rsob.150062.

Crowell, H. L., Soneson, C., Germain, P.-L., Calini, D., Collin, L., Raposo, C., Malhotra, D. and Robinson, M. D. (2020). Muscat detects subpopulation-specific state transitions from multi-sample multi-condition single-cell transcriptomics data. Nature Communications 11: 6077, doi: 10.1038/s41467-020-19894-4.

Cuomo, A. S., Heinen, T., Vagiaki, D., Horta, D., Marioni, J. C. and Stegle, O. (2022). CellRegMap: a statistical framework for mapping context-specific regulatory variants using scRNA-seq. Molecular Systems Biology 18: e10663, doi: 10.15252/msb.202110663.

Edelman, A., Arias, T. A. and Smith, S. T. (1998). The geometry of algorithms with orthogonality constraints. SIAM Journal on Matrix Analysis and Applications 20: 303–353, doi: 10.1137/S0895479895290954.

Gabriel, K. R. (1971). The biplot graphic display of matrices with application to principal component analysis. Biometrika 58: 453–467, doi: 10.2307/2334381.

Goldfarbmuren, K. C., Jackson, N. D., Sajuthi, S. P., Dyjack, N., Li, K. S., Rios, C. L., Plender, E. G., Montgomery, M. T., Everman, J. L., Bratcher, P. E. et al. (2020). Dissecting the cellular specificity of smoking effects and reconstructing lineages in the human airway epithelium. Nature Communications 11: 2485, doi: 10.1038/s41467-020-16239-z.

Gretton, A., Borgwardt, K. M., Rasch, M. J., Schölkopf, B. and Smola, A. (2012). A kernel two-sample test. The Journal of Machine Learning Research 13: 723–773.

Haghverdi, L., Lun, A. T., Morgan, M. D. and Marioni, J. C. (2018). Batch effects in single-cell RNA-sequencing data are corrected by matching mutual nearest neighbors. Nature Biotechnology 36: 421–427, doi: 10.1038/nbt.4091.

Hrvatin, S., Hochbaum, D. R., Nagy, M. A., Cicconet, M., Robertson, K., Cheadle, L., Zilionis, R., Ratner, A., Borges-Monroy, R., Klein, A. M. et al. (2018). Single-cell analysis of experience-dependent transcriptomic states in the mouse visual cortex. Nature Neuroscience 21: 120–129, doi: 10.1038/s41593-017-0029-5.

Jäkel, S., Agirre, E., Mendanha Falcão, A., Van Bruggen, D., Lee, K. W., Knuesel, I., Malhotra, D., Ffrench-Constant, C., Williams, A. and Castelo-Branco, G. (2019). Altered human oligodendrocyte heterogeneity in multiple sclerosis. Nature 566: 543–547, doi: 10.1038/s41586-019-0903-2.

Kang, H. M., Subramaniam, M., Targ, S., Nguyen, M., Maliskova, L., McCarthy, E., Wan, E., Wong, S., Byrnes, L., Lanata, C. M. et al. (2018). Multiplexed droplet single-cell RNA-sequencing using natural genetic variation. Nature Biotechnology 36: 89–94, doi: 10.1038/nbt.4042.

Kim, H. J., Adluru, N., Collins, M. D., Chung, M. K., Bendlin, B. B., Johnson, S. C., Davidson, R. J. and Singh, V. (2014). Multivariate general linear models (MGLM) on Riemannian manifolds with applications to statistical analysis of diffusion weighted images. In Proceedings of the IEEE Conference on Computer Vision and Pattern Recognition, 2705–2712, doi: 10.1109/CVPR.2014.352.

Kim, S., Kim, E., Hitomi, M., Oh, S., Jin, X., Jeon, H., Beck, S., Kim, J., Park, C., Chang, S. et al. (2015). The lim-only transcription factor LMO2 determines tumorigenic and angiogenic traits in glioma stem cells. Cell Death & Differentiation 22: 1517–1525, doi: 10.1038/cdd.2015.7.

Korsunsky, I., Millard, N., Fan, J., Slowikowski, K., Zhang, F., Wei, K., Baglaenko, Y., Brenner, M., Loh, P.-r. and Raychaudhuri, S. (2019). Fast, sensitive and accurate integration of single-cell data with Harmony. Nature Methods 16: 1289–1296, doi: 10.1038/s41592-019-0619-0.

Lähnemann, D., Köster, J., Szczurek, E., Mc-Carthy, D. J., Hicks, S. C., Robinson, M. D., Vallejos, C. A., Campbell, K. R., Beerenwinkel, N., Mahfouz, A. et al. (2020). Eleven grand challenges in single-cell data science. Genome Biology 21: 1–35, doi: 10.1186/s13059-020-1926-6.

Law, C. W., Zeglinski, K., Dong, X., Alhamdoosh, M., Smyth, G. K. and Ritchie, M. E. (2020). A guide to creating design matrices for gene expression experiments. F1000Research 9: 1444, doi: 10.12688/f1000research.27893.1.

Li, G., Yang, D., Nobel, A. B. and Shen, H. (2016). Supervised singular value decomposition and its asymptotic properties. Journal of Multivariate Analysis 146: 7–17, doi: 10.1016/j.jmva.2015.02.016.

Lopez, R., Regier, J., Cole, M. B., Jordan, M. I. and Yosef, N. (2018). Deep generative modeling for single-cell transcriptomics. Nature Methods 15: 1053–1058, doi: 10.1038/s41592-018-0229-2.

Lotfollahi, M., Klimovskaia Susmelj, A., De Donno, C., Hetzel, L., Ji, Y., Ibarra, I. L., Srivatsan, S. R., Naghipourfar, M., Daza, R. M., Martin, B. et al. (2023). Predicting cellular responses to complex perturbations in high-throughput screens. Molecular Systems Biology: e11517doi: 10.15252/msb.202211517.

Lotfollahi, M., Wolf, F. A. and Theis, F. J. (2019). scGen predicts single-cell perturbation responses. Nature Methods 16: 715–721, doi: 10.1038/s41592-019-0494-8.

Love, M. I., Huber, W. and Anders, S. (2014). Moderated estimation of fold change and dispersion for RNA-seq data with DESeq2. Genome Biology 15: 1–21, doi: 10.1186/s13059-014-0550-8.

Madrigal, A., Lu, T., Soto, L. M. and Najafabadi, H. S. (2023). A unified model for interpretable latent embedding of multi-sample, multi-condition single-cell data. bioRxiv doi: 10.1101/2023.08.15.553327.

McInnes, L., Healy, J. and Melville, J. (2018). UMAP: Uniform manifold approximation and projection for dimension reduction. arXiv doi: 10.48550/arXiv.1802.03426.

Missarova, A., Dann, E., Rosen, L., Satija, R. and Marioni, J. (2024). Leveraging neigh-borhood representations of single-cell data to achieve sensitive de testing with milode. Genome Biology doi: 10.1186/s13059-024-03334-3.

Morishima, Y., Gotoh, Y., Zieg, J., Barrett, T., Takano, H., Flavell, R., Davis, R. J., Shirasaki, Y. and Greenberg, M. E. (2001). β-amyloid induces neuronal apoptosis via a mechanism that involves the c-Jun N-terminal kinase pathway and the induction of Fas ligand. Journal of Neuroscience 21: 7551–7560, doi: 10.1523/JNEUROSCI.21-19-07551.2001.

Nathan, A., Asgari, S., Ishigaki, K., Valencia, C., Amariuta, T., Luo, Y., Beynor, J. I., Baglaenko, Y., Suliman, S., Price, A. L. et al. (2022). Single-cell eQTL models reveal dynamic T cell state dependence of disease loci. Nature 606: 120–128, doi: 10.1038/s41586-022-04713-1.

Neufeld, A., Gao, L. L., Popp, J., Battle, A. and Witten, D. (2022). Inference after latent variable estimation for single-cell RNA sequencing data. Biostatistics doi: 10.1093/biostatistic-s/kxac047.

Pijuan-Sala, B., Griffiths, J. A., Guibentif, C., Hiscock, T. W., Jawaid, W., Calero-Nieto, F. J., Mulas, C., Ibarra-Soria, X., Tyser, R. C., Ho, D. L. L. et al. (2019). A single-cell molecular map of mouse gastrulation and early organogenesis. Nature 566: 490–495, doi: 10.1038/s41586-019-0933-9.

Reyfman, P. A., Walter, J. M., Joshi, N., Anekalla, K. R., McQuattie-Pimentel, A. C., Chiu, S., Fernandez, R., Akbarpour, M., Chen, C.-I., Ren, Z. et al. (2019). Single-cell transcriptomic analysis of human lung provides insights into the pathobiology of pulmonary fibrosis. American Journal of Respiratory and Critical Care Medicine 199: 1517–1536, doi: 10.1164/rccm.201712-2410OC.

Robinson, M. D., McCarthy, D. J. and Smyth, G. K. (2010). edgeR: a Bioconductor package for differential expression analysis of digital gene expression data. Bioinformatics 26: 139–140, doi: 10.1093/bioinformatics/btp616.

Sathyamurthy, A., Johnson, K. R., Matson, K. J., Dobrott, C. I., Li, L., Ryba, A. R., Bergman, T. B., Kelly, M. C., Kelley, M. W. and Levine, A. J. (2018). Massively parallel single nucleus transcriptional profiling defines spinal cord neurons and their activity during behavior. Cell Reports 22: 2216–2225, doi: 10.1016/j.celrep.2018.02.003.

Saunders, L. M., Srivatsan, S. R., Duran, M., Dorrity, M. W., Ewing, B., Linbo, T. H., Shendure, J., Raible, D. W., Moens, C. B., Kimelman, D. and Trapnell, C. (2023). Embryo-scale reverse genetics at single-cell resolution. Nature 623: 782–791, doi: 10.1038/s41586-023-06720-2.

Skinnider, M. A., Squair, J. W., Kathe, C., Anderson, M. A., Gautier, M., Matson, K. J., Milano, M., Hutson, T. H., Barraud, Q., Phillips, A. A. et al. (2021). Cell type prioritization in single-cell data. Nature Biotechnology 39: 30–34, doi: 10.1038/s41587-020-0605-1.

Smyth, G., Hu, Y., Ritchie, M., Silver, J., Wettenhall, J., McCarthy, D., Wu, D., Shi, W., Phipson, B., Lun, A., Thorne, N., Oshlack, A., Graaf, C. de, Chen, Y., Langaas, M., Ferkingstad, E., Davy, M., Pepin, F. and Choi, D. (2023). limma: linear models for microarray and RNA-seq data. doi: 10.18129/B9.bioc.limma, User guide (v3.58.1) chapter 9.6.2.

Smyth, G. K. (2004). Linear models and empirical bayes methods for assessing differential expression in microarray experiments. Statistical Applications in Genetics and Molecular Biology 3, doi: 10.2202/1544-6115.1027.

Wold, S., Sjöström, M. and Eriksson, L. (2001). PLS-regression: a basic tool of chemometrics. Chemometrics and Intelligent Laboratory Systems 58: 109–130, doi: 10.1016/S0169-7439(01)00155-1.

Yao, Z., Van Velthoven, C. T., Nguyen, T. N., Goldy, J., Sedeno-Cortes, A. E., Baftizadeh, F., Bertagnolli, D., Casper, T., Chiang, M., Crichton, K. et al. (2021). A taxonomy of transcriptomic cell types across the isocortex and hippocampal formation. Cell 184: 3222–3241, doi: 10.1016/j.cell.2021.04.021.

Zhao, W., Dovas, A., Spinazzi, E. F., Levitin, H. M., Banu, M. A., Upadhyayula, P., Sudhakar, T., Marie, T., Otten, M. L., Sisti, M. B. et al. (2021). Deconvolution of cell type-specific drug responses in human tumor tissue with single-cell RNA-seq. Genome Medicine 13: 82, doi: 10.1186/s13073-021-00894-y.

Zhou, Y., Luo, K., Liang, L., Chen, M. and He, X. (2023). A new Bayesian factor analysis method improves detection of genes and biological processes affected by perturbations in single-cell CRISPR screening. Nature Methods 20: 1693–1703, doi: 10.1038/s41592-023-02017-4.

